# *KAT6A* mutations drive transcriptional dysregulation of cell cycle and Autism risk genes in an Arboleda-Tham Syndrome cerebral organoid model

**DOI:** 10.1101/2023.06.17.545322

**Authors:** Aileen A. Nava, Connor T. Jops, Celine K. Vuong, Samantha L. Niles-Jensen, Leroy Bondhus, Cameron J. Ong, Luis de la Torre-Ubieta, Michael J. Gandal, Valerie A. Arboleda

## Abstract

Arboleda-Tham Syndrome (ARTHS, OMIM#616268) is a rare neurodevelopmental disorder caused by *de novo* mutations in *KAT6A*. Individuals with ARTHS typically exhibit varying degrees of intellectual disability, speech and language deficits and clinical manifestations across multiple systems that lead to abnormal: vision, craniofacial features, cardiac morphology, and gastrointestinal function. To gain insight into the potential neuropathological mechanisms underlying ARTHS, we investigate how *KAT6A* mutations disrupt *in vitro* brain development using induced pluripotent stem cells (iPSCs) and cerebral organoids (COs) derived from ARTHS patients harboring *KAT6A* nonsense mutations. In this study, we conducted comprehensive transcriptomic profiling by performing time-course experiments and generating short-read and long-read RNA sequencing (RNA-seq) data from undifferentiated iPSCs and COs at 15 and 25 days of neural differentiation. Our analysis revealed abnormal expression of 235 genes in ARTHS across all three timepoints examined. Notably, we observed persistent dysregulation of genes such as *CTSF*, *ZNF229*, *PCDHB12*, and *PAK3*. Additionally, we found a consistent enrichment of *PTBP1*-target genes among the upregulated genes in ARTHS at all three stages assessed by RNA-seq. During neural differentiation, we identified 980 genes that consistently display aberrant transcription in ARTHS at both CO stages. These genes are enriched for genes involved in cell fate determination through modulation of cell-cycle dynamics (e.g. *E2F* family) and cell-adhesion molecules (e.g. *PCDH* genes). Our findings indicate that ARTHS COs exhibit slower downregulation of pluripotency and cell cycle genes compared to controls and that this delay led to an overrepresentation of cycling human neural progenitor markers during neural differentiation in ARTHS. Finally, matching the variable neurodevelopment phenotypes in ARTHS, we discovered that the aberrantly expressed genes in ARTHS are enriched for genes associated with Autism Spectrum Disorder and Epilepsy, with a subset showing isoform-specific dysregulation. Strikingly, the same *PTBP1-*target genes were enriched amongst the genes that display differential isoform usage in ARTHS. For the first time, we demonstrate that *KAT6A* mutations lead to a delay in repressing pluripotency and cell cycle genes during neural differentiation, suggesting that prolonged activation of these gene networks disrupts the temporal dynamics of human brain development in ARTHS.

## Introduction

Arboleda-Tham Syndrome (ARTHS, OMIM#616268), also known as ‘KAT6A Syndrome’, is a rare monogenic neurodevelopmental disorder (NDD). First described in 2015 ^1,2,3^, it’s caused by autosomal dominant mutations in the *Lysine (K) acetyltransferase* 6A (*KAT6A)* gene located on chromosome 8p11.21 and encodes a 2004 amino acid protein that modulates the epigenome^4^ —making it part of a large group of proteins known as epigenes ^5,6^. A majority of individuals with ARTHS carry either protein-truncating mutations in *KAT6A* ^7–15^ or missense mutations localized to highly conserved regions of the *KAT6A* gene ^16–21^.

ARTHS patients present with a spectrum of disease-related phenotypes that originate from disruptions of early developmental processes, a common feature in congenital syndromes caused by germline mutations in epigenes ^4^. All ARTHS patients display global developmental delay, intellectual disability, speech and language deficits. However, more variable features include: feeding difficulties, hypotonia, vision problems, gastrointestinal issues, sleep disturbances and craniofacial and cardiac malformations ^8,11^. Notably, seizure disorders ^8,11^ and Autism Spectrum Disorder (ASD) co-occur in approximately 30% of ARTHS patients ^8,11,19^. Studies have shown that ARTHS patients with “late mutations” (protein-truncating mutations in exons 16-17 of *KAT6A*) typically experience more severe symptoms than those with “early mutations” in *KAT6A* (protein-truncating mutations in exons 1-15)^11^. The rapid clinical characterization of these patients has illuminated a complex picture of the spectrum of clinical manifestations. However, the molecular mechanisms driving these clinical phenotypes remain poorly characterized.

*KAT6A* is an evolutionarily conserved lysine (K) acetyltransferase (KAT) ^22,23^ that catalyzes the transfer of acetyl groups to histone tails to regulate gene expression ^4,24^. Model organism studies have shown regulation of lysine acetylation is required to ensure neuroprogenitor cell (NPC) proliferation and enable the differentiation of neural lineages (e.g. neurons, astrocytes, and oligodendrocytes) ^25^ during brain development. One limitation of studies in model organisms is that they do not account for the subtle, but meaningful, species-to-species variability. Studies focused on human cells, with endogenous KAT6A expression, and modeling early developmental time points, are critical to dissect out the earliest perturbations that underlie ARTHS pathophysiology.

Induced pluripotent stem cells (iPSCs) present an effective model for studying human development and possess the capacity to differentiate into almost every cell type found in the human body. The study of neurodevelopmental disorders has been revolutionized by use of patient-derived iPSCs which facilitate study of early neurodevelopmental through *in vitro* modeling using cerebral organoids (CO)^4,26–31,32–34^. COs are miniature three-dimensional (3D) models of the developing brain and they are typically generated through the self-organization and differentiation of iPSCs into various NPC-derived lineages ^35–37^. Cerebral organoids (CO) created from patient-derived iPSCs can link genetic mutations to molecular and cellular defects occurring in early brain development^35–37^. At early stages of CO development, the COs show neural rosette structures and are predominantly composed of primitive cell types, such as like NPCs ^35,36^ and after 30 days increase cellular diversity and the cellular networks resemble more mature brain components. We sought to understand how KAT6A mutations influence the earliest stages of brain development and leveraged bulk assays to characterize the role of KAT6A mutations during ARTHS ND.

In normal ND, neuroepithelial stem/progenitor cells (NPCs) are the elementary units of the brain, which is a multi-faceted tissue composed of complex structures and diverse cell types ^38,39^. The five core biological processes that occur during embryonic development of the brain are: (1) neurulation, (2) proliferation, (3) cell migration, (4) differentiation, and (5) synaptogenesis ^40^. The proper formation of the brain during embryonic development is dependent on the precise spatiotemporal timing of both proliferation and differentiation to generate the appropriate number of NPC-derived lineages in the correct position within the adult brain ^41^.

Critically, NPCs in the developing brain undergo an initial wave of proliferation, before exiting the cell cycle to enable differentiation as they migrate out away from these proliferative regions to achieve their final cell fates ^38^. Importantly, in the developing human brain, cycle cycle progression is transcriptionally linked to early cell fate choices during neurogenesis - demonstrating neural differentiation exists on a ‘transcriptomic continuum’ ^42^. In the developing brain, previous studies have demonstrated that cell cycle dynamics, capacity to differentiate, and the cellular fates of NPCs are co-regulated by several neurogenic transcription factors (TFs) and RNA-binding Proteins (RBPs) like those of the *E2F* family ^43–45^ and *PTB* family ^46–52^, respectively. Generally, TFs and RBPs regulate gene expression via pre-transcriptional and post-transcriptional programs that involve binding to specific DNA motifs ^53,54^ and altering splicing of RNA ^52,55^, respectively. *E2F*s are tissue-specific TFs that regulate the cellular fate of various stem cells through transcriptional control of cell cycle dynamics ^56,57^. While members of the *PTB* family (i.e. *PTBP1* and *PTBP2*) are known to regulate cell fate transition of NPCs through modulation of alternative splicing in neural lineages ^49,58–61^. Neural circuitry is formed shortly after differentiation through synaptogenesis which is the process of synapse formation between neurons within the brain ^40,62^. Protocadherin (*PCDH*) genes encode cell-surface adhesion molecules that are prominently expressed in the central nervous system—where they function as signaling receptors to modulate the establishment of neural circuitry in the developing brain through cell-to-cell adhesion ^63–66^ and are linked to common NDDs like Autism Spectrum Disorder (ASD, OMIM#209850) and Down Syndrome ^67–69^. Multiple transcriptional and cell-surface molecules regulate cell fate and when disrupted lead to neurodevelopmental disorders.

We have, for the first time, delineated the altered molecular landscape in ARTHS during *in vitro* brain development by deeply profiling iPSCs and cerebral organoids (COs) derived from ARTHS patients. Using a time-course transcriptomic analysis across three timepoints of early CO differentiation, we identified dysregulated molecular networks and biological pathways driven by pathogenic *KAT6A* mutations. Our study reveals pathogenic *KAT6A* mutations delay the repression of key pluripotency and cell cycle genes during early neural differentiation. This leads to an enrichment of cycling NPC markers that prevent initiation of multiple molecular signals critical for normal neural differentiation. Our results support a key role of RNA-splicing regulator, *PTBP1* and its target genes that are known to regulate cell identity transition of NPCs neural differentiation ^49,59^ and show a delay the upregulation of neural-specific cell-adhesion molecules (e.g. *PCDH* genes) in ARTHS COs. Finally, ARTHS dysregulated genes are significantly enriched for ASD and epilepsy-risk genes, with a subset showing isoform-specific dysregulation. Our work directly assays how *KAT6A* mutations disrupt NPC specification and differentiation in ARTHS and highlights convergent molecular and biological underpinnings of rare and common NDDs.

## Results

### Generation and Verification of ARTHS Induced Pluripotent Stem Cells (iPSCs)

To study early neurodevelopmental events in ARTHS, we generated iPSC lines from peripheral blood mononuclear cells (PBMCs) obtained from two ARTHS patients (**Fig. S1A**). Both patients had a previous clinical genetic diagnosis of ARTHS^2,11^ and have *de novo*, heterozygous *KAT6A* mutations that are predicted to cause premature protein truncation (**Fig. 1A, Table S1**). Common phenotypic features shared between the ARTHS patients include intellectual disability, speech delays, visual defects, strabismus, distinctive facial features, and congenital heart defects. Neither patient was small for their gestational age, nor was diagnosed with microcephaly or seizures. Since the goal of our study was to explore the earliest stages of neural specification that drive the neurodevelopmental delays in ARTHS patients, we leveraged a 3D model of *in vitro* neurodevelopment in which iPSCs can differentiate into all cell types, including early brain precursor cells. As controls, we obtained sex- and age-matched iPSCs generated at the same Biomanufacturing Center that generated our ARTHS lines (**Table S2**) and were confirmed to have no clinical phenotypes.

**Fig 1:**
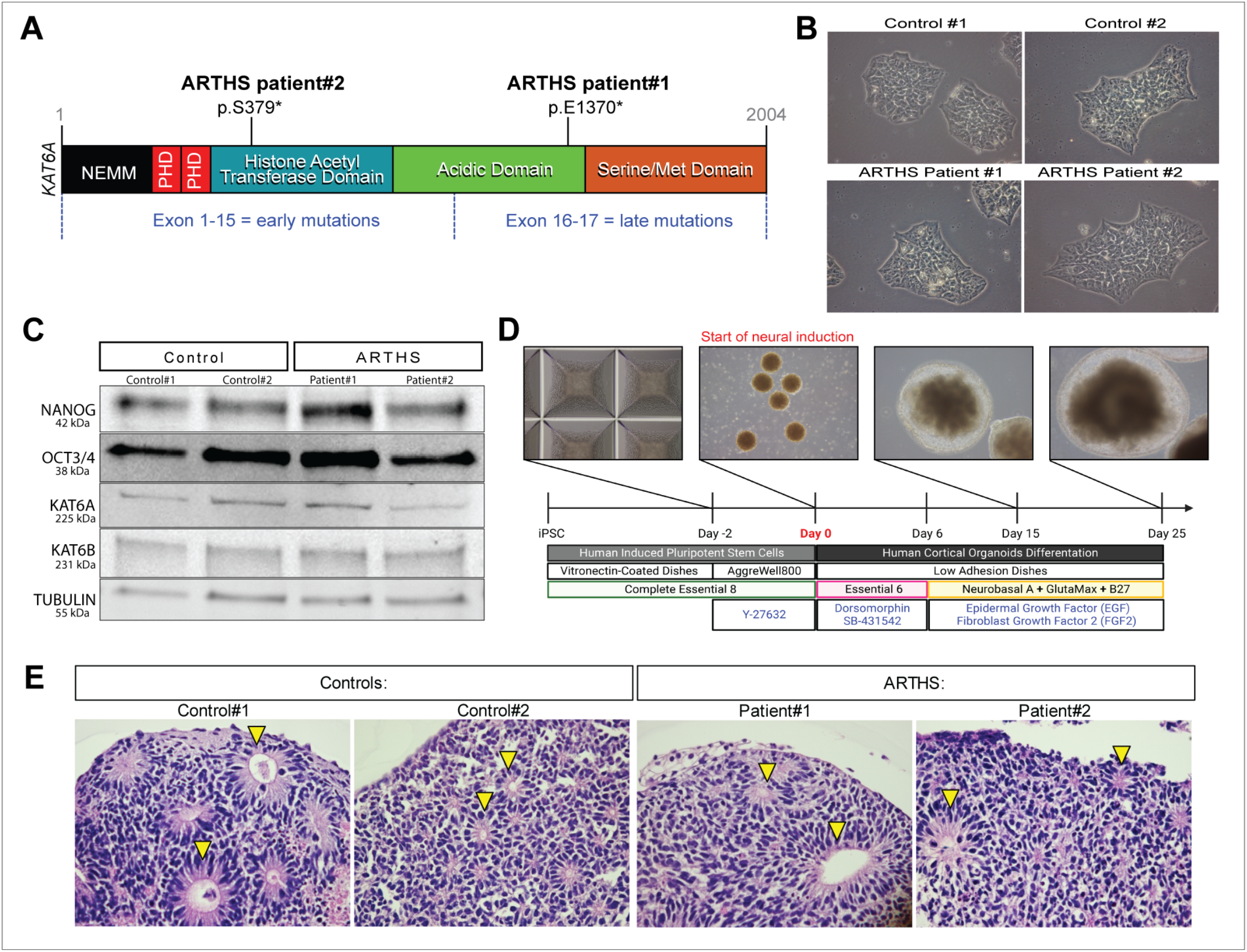
Generation and characterization of iPSCs and COs derived from ARTHS patients. **(A)** Location of ARTHS patient mutations along the *KAT6A* gene: ARTHS patient#1 and patient#2 both have heterozygous nonsense *KAT6A* mutations that result in premature truncation codon at residue 1370 and 379, respectively; annotations derived from the following NCBI reference sequences: NG_042093.1, NM_006766.5, NP_006757.2. **(B)** Brightfield images of the four iPSC lines used in this study reveal they exhibit the expected iPSC morphology. **(C)** Western blot analysis of nuclear protein from the four iPSC lines used in this study; protein quantification of this western blot, which available in supplementary material, shows control and ARTHS iPSC lines express indistinguishable amounts of NANOG, OCT3/4, and KAT6B protein. ARTHS iPSC lines express significantly (p-value <0.05) less KAT6A protein compared to control iPSC lines. **(D)** Overview of the cerebral organoid differentiation process used in this study; RNA sequencing was performed at three points along this timeline: the “IPSC stage”, day 15 (CO15), and day 25 (CO25). **(E)** At day 25, COs were assessed for the formation of neural rosettes via hematoxylin & eosin staining; examples of neural rosettes are denoted with a yellow triangle and demonstrate successful differentiation of iPSCs into early brain-like tissue.

We confirmed that all iPSC lines retained the cellular morphology of undifferentiated iPSCs (**Fig. 1B**), matched *KAT6A* genotype (**Fig. S1B**) and expressed key pluripotency markers *OCT3/4* (also known as *POU5F1*), *NANOG*, *SOX2*, *TRA-1-60*, *TRA-1-81*, and *SSEA4* (**Fig. S1C** and *data not shown*) Immunofluorescence and western blot of iPSC for *NANOG* and *OCT3/4* show no differences in expression levels or localization between ARTHS or controls (**Fig. 1C, Fig. S2-S3**). However, KAT6A protein is significantly decreased in ARTHS compared to control iPSCs (**Fig. 1C, Fig. S2**), suggesting that truncating mutations in *KAT6A* lead to haploinsufficiency in ARTHS patients. No significant changes in the protein expression of KAT6B protein (a paralog of *KAT6A*) were observed in ARTHS iPSCs (**Fig. 1C, Fig. S2**). Taken together, we show that high-quality iPSCs were generated from ARTHS patients and that *KAT6A* mutations lead to decreased protein expression but do not alter iPSC pluripotency markers.

### Generation and Characterization of Cerebral Organoids (COs) created from ARTHS-derived iPSCs

To investigate the effects of nonsense *KAT6A* mutations on *in vitro* human ND, we created cerebral organoids (COs) from ARTHS and control iPSC lines using a reproducible differentiation protocol (**Fig. 1D**) ^70^. After 15 days of CO differentiation (CO15), we observed formation of radial lumen reminiscent of neural-rosette-like structures through brightfield imaging (**Fig. 1E**). By 25 days of differentiation (CO25), organoids maintained the neural rosette structures and expressed PAX6 and NESTIN protein (**Fig. S4A**). Since pathogenic *KAT6A* mutations are associated with microcephaly in ARTHS, we asked if COs derived from ARTHS iPSCs developed smaller organoid structures than control iPSCs. We used QuPath software ^71^ to quantify the average size of ARTHS and control COs at day 25 of differentiation (**Fig. 1E**) but did not detect significant differences in CO size (**Fig. S4B**).

### Pathogenic mutations in KAT6A drive global transcriptomic dysregulation across all stages of CO differentiation

*KAT6A* is an enzyme with histone acetyltransferase activity that deposits acetyl groups onto lysine residues present on histone tails, modulates chromatin organization and accessibility ^22,72–75^ and therefore can limit the availability of the transcriptional machinery that drives gene expression ^76,77^. *In vivo* studies have established that *KAT6A* is required for normal development of multiple organ systems ^78–91^. Since KAT6A is required for transcriptional regulation and normal embryonic development, we hypothesized that pathogenic *KAT6A* mutations will result in transcriptional dysregulation in an *in vitro* model of neurodevelopment.

To assess the neurodevelopmental trajectory of our *in vitro* model, we conducted transcriptome profiling at 3 stages along the CO differentiation processes (i.e. iPSC, CO15, CO25; **Fig. 1D**). Specifically, we performed high-coverage, short-read RNA sequencing (RNA-seq) with 3 replicates from independent wells of CO generation for each patient sample for a total of 36 RNA-seq libraries. At each of the three time points (n=3) along CO differentiation, we assayed ARTHS patients (n=2) and controls (n=2) and each biological individual was sampled in triplicate during the differentiation protocol (n=3). All RNA-seq libraries achieved on average 73.7 million paired-end reads (**Table S3-S5**) and principal component analysis revealed that the samples separated along PC1 based on their progression through *in vitro* CO differentiation (**Fig. S5**) ^70^.

Differential gene expression analysis at each time point (i.e. IPSC, CO15, CO25) using DESeq2 quantified the gene expression fold change (FC) between ARTHS to control samples ^92,93^. To identify how pathogenic *KAT6A* mutations affect the transcriptome of each cell lineage during CO differentiation—for each individual list of sigDE genes, we performed gene ontology (GO) enrichment analysis using ClusterProfiler ^94^. At the IPSC stage, a total of 1931 genes were sigDE in ARTHS relative to unaffected controls (Benjamini-Hochberg adjusted p-value (P_adj_) < 0.05, **Fig. 2A**, **Fig. S6**, **Table S6**). Upon classifying the IPSC sigDE, we found this list mainly consisted of ‘protein coding’ (75.30%) and ‘lncRNA’ (15.74%) genes—with most genes showing an up-regulation in expression (57.22%,1105/1931). IPSC sigDE genes were significantly enriched for a total of 57 GO terms (P_adj_ < 0.05, **Table S7**): 10 biological processes (BPs), 38 cellular components (CCs), and 9 molecular functions (MFs). At the IPSC stage, genes that were sigDE in ARTHS are most significantly overrepresented (P_adj_ < 0.05, **Fig. S6C**) in BP GO terms related to the development and morphogenesis of cardiac tissues, in addition to the development of neuronal structures. The IPSC sigDE genes that constitute the top 5 BP GO terms show moderate interconnectedness, highlighting the pleiotropic-nature of many developmental genes (**Fig. S7**).

**Fig 2:**
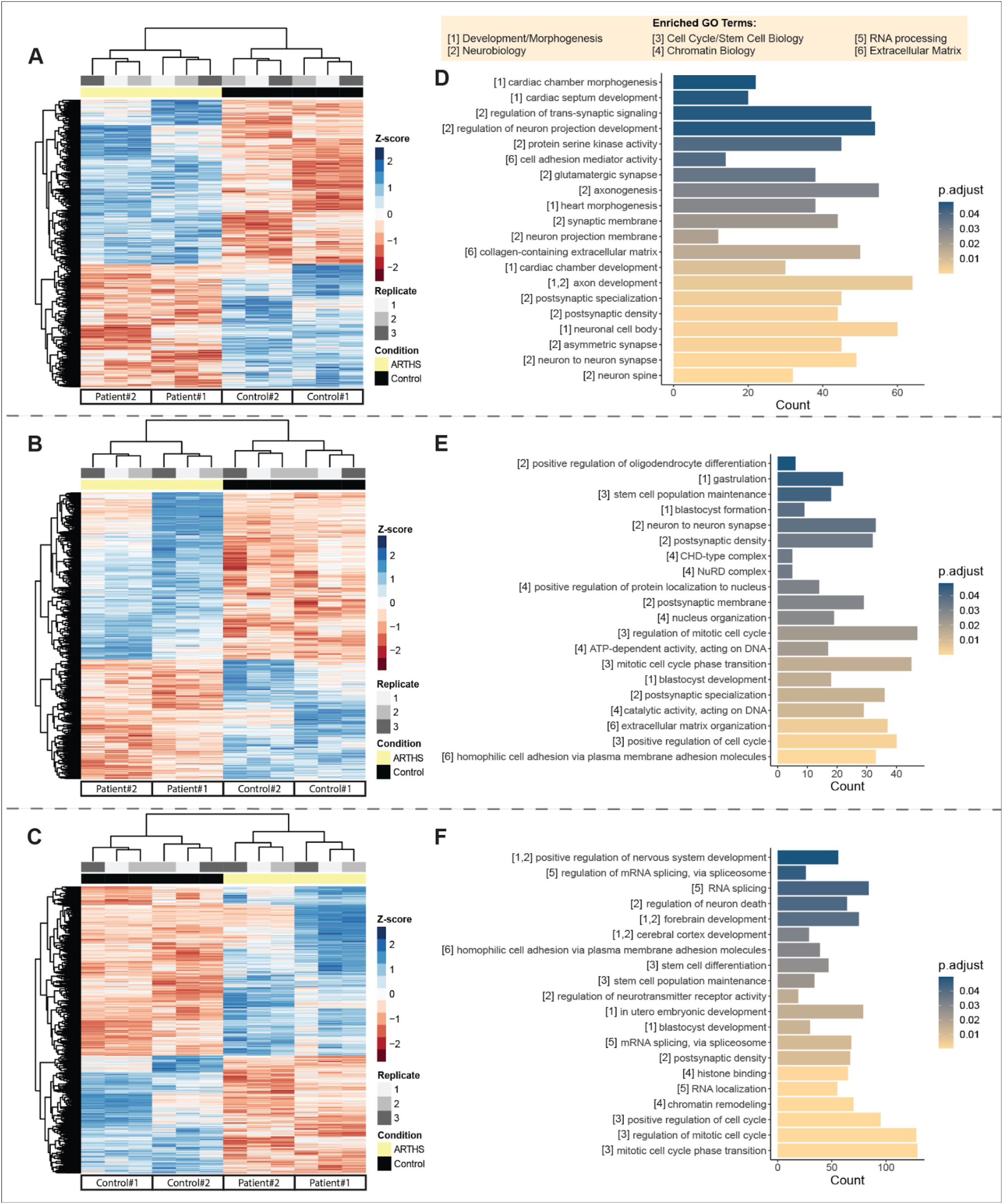
Pathogenic *KAT6A* mutations cause transcriptomic perturbations during CO differentiation. Unsupervised hierarchical clustering of all significantly differentially expressed genes in ARTHS identified significant changes in gene expression at the (A) iPSCs (n= 1,931 sigDEGs), (B) cerebral organoids after 15 days (n= 1,673 sigDEGs), and (C) after 25 days of neural induction (n= 4,088 sigDEGs) (P_adj_ < 0.05). Heat maps display the DESeq2 normalized gene expression Z-scores for all sigDEGs at each timepoint. Gene ontology (GO) enrichment analysis of all sigDEGs in ARTHS (D) iPSCs (n= 57 GO terms), (E) cerebral organoids after 15 days (n= 132 GO terms), and (F) after 25 days of neural induction (n= 390 GO terms) identified significant overrepresentation of 6 core aspects of embryogenesis (P_adj_ < 0.05) amongst all the genes found to be significantly transcriptionally dysregulated in ARTHS relative to unaffected controls: [1] tissue development/morphogenesis, [2] neurobiology, [3] cell cycle/stem cell biology, [4] chromatin biology, [5] RNA processing, and [6] extracellular matrix organization.

Fifteen days after the start of neural induction, at the CO15 stage, we identified 1673 genes that are sigDE in ARTHS compared to controls (P_adj_ < 0.05, **Fig. 2B**, **Fig. S8**, **Table S8**). CO15 sigDE genes were primarily classified as ‘protein coding’ (66.71%) and ‘lncRNA’ (20.44%)—with most genes showing an up-regulation in expression (58.28%, 975/1673). This list of CO15 sigDE genes was significantly enriched for 132 GO terms (P_adj_ < 0.05, **Table S9**): 101 BPs, 27 CCs, and 4 MFs. As shown in **Fig. S8C**, genes that were sigDE in ARTHS at the CO15 stage were most significantly enriched (P_adj_ < 0.05) in BP GO terms related to: cell to cell adhesion, extracellular matrix organization, blastocyst development, and dynamic of DNA replication and cell division. On day 25 of CO differentiation, we identified 4088 genes that were sigDE in ARTHS relative to controls (P_adj_ < 0.05, **Fig. 2C**, **Fig. S9**, **Table S10**). The CO25 sigDE genes were predominantly ‘protein coding’ (70.30%) and ‘lncRNA’ (18.35%) genes—with most genes showing an up-regulation in expression (58.98%, 2411/4088). Also, genes that were sigDE in ARTHS at the CO25 stage were significantly enriched for 390 GO terms (P_adj_ < 0.05, **Table S11**): 292 BPs, 65 CCs, and 33 MFs. While the CO25 sigDE genes were most significantly enriched (P_adj_ < 0.05) for GO terms related to the following BPs: chromatin organization, DNA replication, and cell division (**Fig. S9C**). Lastly, all the genes which were transcriptionally dysregulated in ARTHS patient samples across all three stages of the CO differentiation protocol - were enriched for GO terms related to 6 broad categories (**Fig. 2D-F**): development/morphogenesis, neurobiology, cell cycle/stem cell biology, chromatin biology, RNA processing, and extracellular matrix.

### Lineage-independent dysregulated genes are enriched for genes involved in neuronal differentiation

Transcript expression of *KAT6A* and *KAT6B* during our *in vitro* ND protocol differentiation increased in both ARTHS and control samples (**Fig. 3A** and **Fig. S10**). However, we noted that ARTHS samples expressed less KAT6A RNA compared to controls, with this trend reaching statistical significance at day 25 of differentiation (Log_2_FC = −0.280, P_adj_= 4.09×10^-5^) (**Fig. 3A**). No significant differences in *KAT6B,* a close homolog of *KAT6A,* RNA levels were detected (**Fig. S10**).

**Fig 3:**
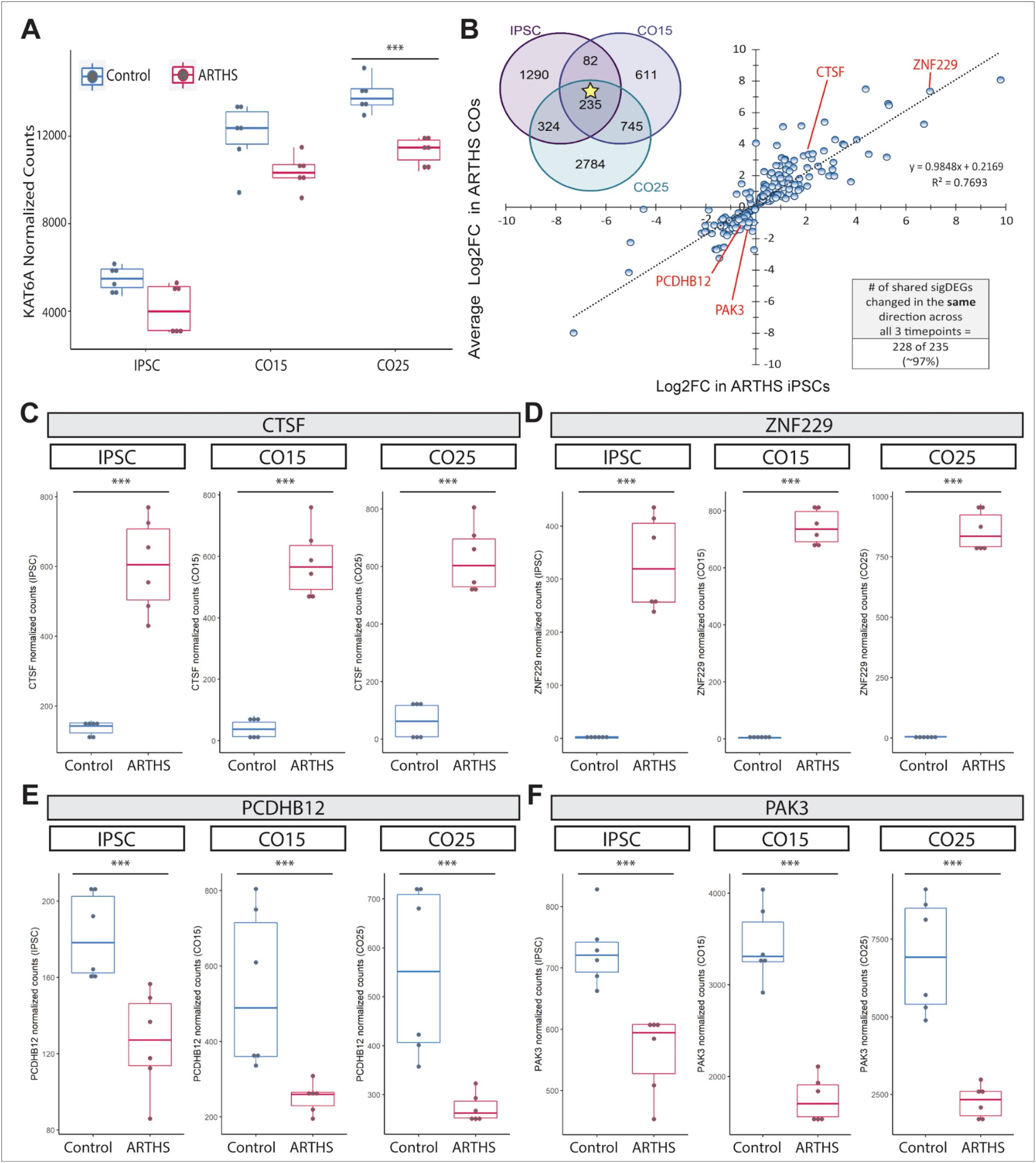
Transcriptionally dysregulated and lineage independent genes disrupt neural specification networks. (A) The normalized gene expression of *KAT6A* is plotted across all 3 stages of CO differentiation and a trend was observed where ARTHS biosamples generally expressed lower amounts of KAT6A transcripts across time, with this trend reaching significance at CO25 (KAT6A Log_2_FC in ARTHS at CO25= −0.28, Padj= 4.09×10^-5). (B) We identified 235 genes that are transcriptionally dysregulated in lineage independent manner across all 3 timepoints in ARTHS-patient biosamples. The Log_2_FC of these 235 sigDE genes is highly correlated across the IPSC and COs stages as (R² of 0.7693) and 97% of genes change in the same direction. Here we highlight the top four neurodevelopmental-related genes, (C) CTSF, (D) ZNF229, (E) PCDHB12, (F) PAK3, that show significant transcriptional perturbation (Padj <0.05) across all 3 timepoints of CO differentiation in ARTHS-cells.

Across all three timepoints, we identified 235 sigDE genes (235/6071 or 3.87%) that are consistently dysregulated in ARTHS (**Fig. 3B**, **Tables S12**) and appear to be lineage independent in their dysregulation by KAT6A. To investigate the convergent transcriptomic dysregulation caused by pathogenic *KAT6A* mutations, we then correlated the relative changes in gene expression (i.e. ARTHS/control) for this conserved list of 235 sigDE genes that were consistently altered in ARTHS at all three cell stages of CO differentiation and found a strong positive correlation (R^2^=0.7693) between the log_2_FC calculate at the IPSC and CO stages (**Fig. 3B**) and 228 out of 235 conserved lineage-independent sigDE genes (97.02%) were transcriptionally dysregulated in the same direction across all cell stages (IPSC, CO15, CO25).

This conserved list of 235 lineage-independent sigDE genes contained several significantly upregulated and downregulated genes in ARTHS patient biosamples across all three stages of the CO differentiation. In **figures 3C-F**, we highlight two neuronal differentiation related genes from each gene-regulation paradigm. We show that *CTSF* was significantly elevated in ARTHS at the IPSC (Log_2_FC = 2.11, P_adj_= 5.07×10^-38^), CO15 (Log_2_FC = 3.79, P_adj_= 3.27×10^-14^), and CO25 (Log_2_FC = 2.92, P_adj_= 3.44×10^-04^) stages relative to unaffected controls (**Fig. 3C**). We also find that a zinc finger protein (ZFP), *ZNF229*, is significantly overexpressed in ARTHS relative to controls at the IPSC stage (Log_2_FC = 6.96, P_adj_= 2.78×10^-59^), CO15 stage (Log_2_FC = 7.16, P_adj_= 5.07×10^-138^), and CO25 stage (Log_2_FC = 7.56, P_adj_= 1.92×10^-137^) (**Fig. 3D**). On average, the expression of *ZNF229* was upregulated by 150-fold in ARTHS compared to unaffected controls, making it the most transcriptionally dysregulated gene in our srRNA-seq data. Next, we looked at genes which were consistently downregulated in ARTHS across all stages of CO differentiation. Strikingly, we found *PCDHB12* was significantly repressed in ARTHS at the iPSC (Log_2_FC = −0.44, P_adj_= 8.80×10^-03^), CO15 (Log_2_FC = −0.97, P_adj_= 1.37×10^-03^), and CO25 (Log_2_FC = −0.93, P_adj_= 2.86×10^-05^) stages relative to controls (**Fig. 3E**). We also identified that *PAK3*, a key regulator of synaptic plasticity that causes a form of non-syndromic X-linked intellectual disability, was significantly depleted in ARTHS at the IPSC (Log_2_FC = −0.33, P_adj_= 2.03×10^-03^), CO15 (Log_2_FC = −0.94, P_adj_= 6.18×10^-11^), and CO25 (Log_2_FC = −1.56, P_adj_= 2.05×10^-12^) stages relative to controls (**Fig. 3F**). Taken together, this data shows that pathogenic truncating *KAT6A* mutations in ARTHS cause significant transcriptional perturbations that are preserved across neural and non-neural cell lineages.

At both the CO15 and CO25 stages, we discovered that 30 and 32 *PCDH* genes, respectively, exhibited significant transcriptional dysregulation in ARTHS—with most *PCDH* genes (27/30 and 30/32) showing a significant reduction in transcription compared to unaffected control COs (**Tables S8 and S10**). Finally, these transcriptionally dysregulated *PCDH* genes constitute a large proportion of the top two BP GO terms that are significantly overrepresented amongst the genes found to be sigDE in ARTHS at the CO15 stage (**Table S9**, **Fig. S11**). *PCDH* genes are known to play an integral role in normal brain development, making these key candidate genes underlying ARTHS neuropathology.

### KAT6A mutations disrupt cell-cycle genes during early neurodevelopment

To delineate the roles of the genes that are most transcriptionally dysregulated in ARTHS during CO differentiation, we next took differentially expressed genes across CO15 and CO25, to determine if there were specific networks perturbed during *in vitro* neurodevelopment. We identified 980 genes (**Table S13**) that were dysregulated in both CO timepoints compared to controls. ClusterProfiler GO enrichment analysis of these 980 persistently sigDE genes identified 167 GO terms (P_adj_ <0.05, **Table S14**) that were related to two overarching BPs (**Figs. S12-S13**): cell cycle dynamics (e.g. GO:0045787, GO:0044772, GO:0044786) and cell-cell adhesion (e.g. GO:0098742). This suggested that pathogenic *KAT6A* mutations disrupt the cell cycle - neural differentiation transition during early neurodevelopment.

Cell cycle dynamics are known to modulate the differentiation of both pluripotent stem cells ^57,95,96^ and NPCs ^56,97,98^. We hypothesized that specific TFs are one of the factors at the crux of the aberrant transcription observed in ARTHS during CO differentiation since some of the top enrichments identified upon analyzing the genes which were sigDE in ARTHS COs were related to cell fate modulation, DNA-binding, and cell cycle dynamics (**Fig. 2** and **Figs. S8-S9**, **S11**, **S12-S15**). We used HOMER to identify TF binding motifs that were enriched within the promoters of the sigDE genes. We found 11 known and 39 *de novo* DNA-binding motifs (p-value<0.05, **Table S15**). These sigDE in ARTHS COs were significantly enriched for DNA motifs that are bound by neurogenic TFs (**Fig. 4A**) and enriched for gene networks associated with NPC differentiation, pluripotency, and E2F targets ^99–102^ (p-value<0.05, **Table S15**). Among the top 5 motifs enriched in the persistently sigDE genes in ARTHS COs, we found 3 DNA motifs known to be bound by members of the *E2F* family (**Fig. 4A**). In total, 21.84% (214/980) of all sigDE genes contained DNA motifs in their promoter corresponding to the transcription factor *E2F3*, a master regulator of cell cycle genes during neuronal differentiation.

**Fig 4:**
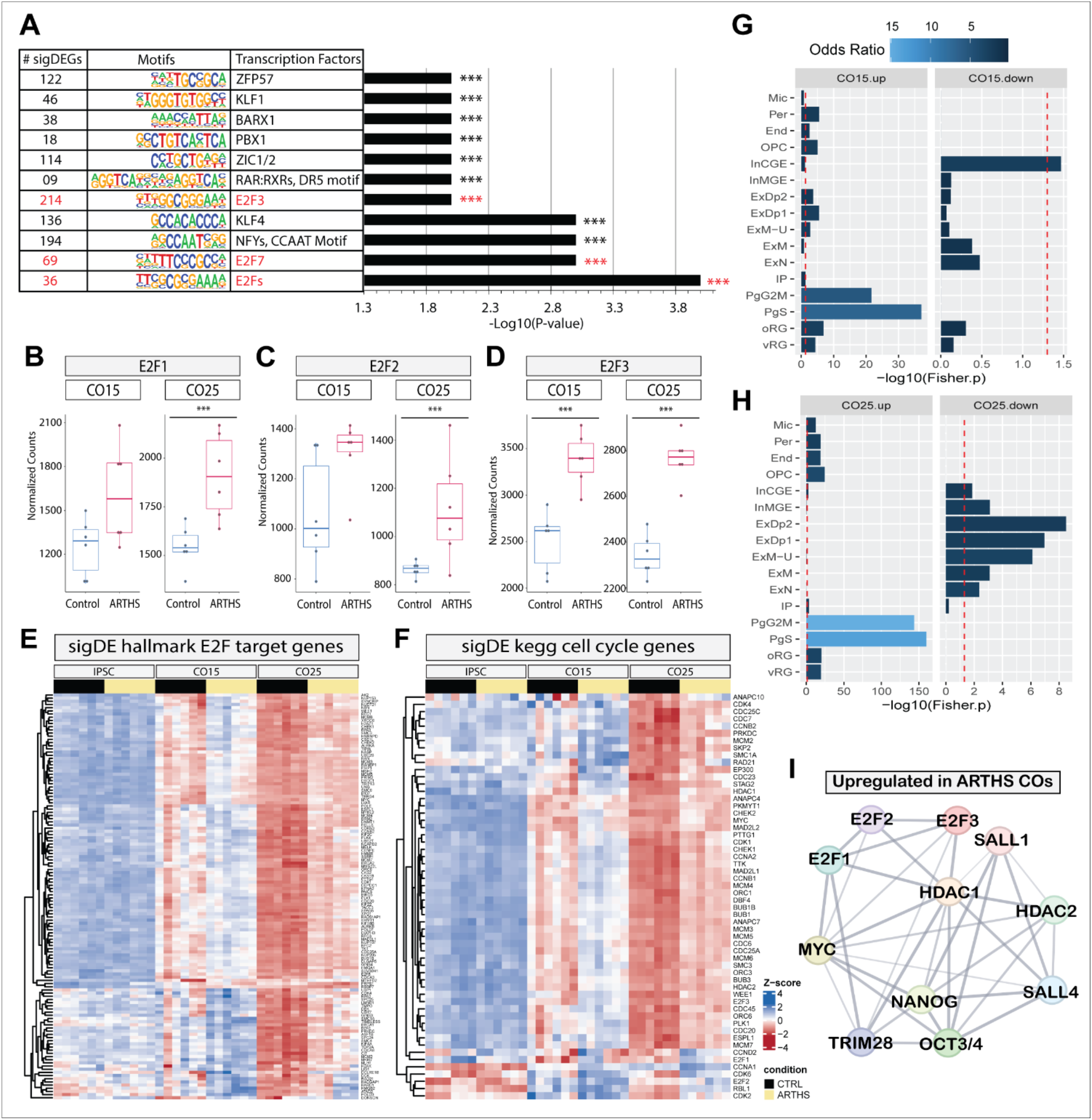
Pathogenic KAT6A mutations disrupt cell-fate transcriptional modules in ARTHS COs. Cumulatively, we identified 4,784 unique genes that are significantly differentially expressed (sigDE) in ARTHS cerebral organoids at either day 15 and day 25 of differentiation (i.e. CO15, CO25) relative to controls (P_adj_ < 0.05). We found 980/4,784 sigDE genes were consistently sigDE in ARTHS at both CO15 and CO25. (A) DNA motif enrichment analysis of the 980 genes which are consistently sigDE in ARTHS COs identified 11 motifs that are significantly enriched (p-value<0.05). All 11 motifs corresponded to neurogenic transcription factors. 3 of 11 transcription factor motifs (27.3%) correspond to proteins within the E2F transcription factor family (red) and 214 of 980 sigDE genes (21.84%) contain motifs corresponding to E2F3, an established master regulator of the cell cycle during neural differentiation. We also observed a significant overexpression of (B) E2F1, (C) E2F2, and (D) E2F3 in ARTHS throughout CO differentiation. Across the differentiation protocol, ARTHS COs showed a significant delay in down-regulating the transcriptional expression of (E) 128 ‘hallmark E2F target’ genes and (F) 56 ‘KEGG cell cycle’ genes relative to control COs. (G-H) Our cell-type enrichment analysis revealed a significant overrepresentation of single-cell markers corresponding to cycling NPCs and different neuronal populations within the developing human fetal brain amongst all the sigDE genes which were either up-regulated (‘up’) or down-regulated (‘down’) in ARTHS COs, respectively (p-value<0.05). Notably, we observed a consistent enrichment of G2/M phase cycling NPCs (ie. ‘PgG2M’) amongst up-regulated genes identified in ARTHS at CO15 (Fisher’s Exact test: OR=5.47, p-value=2.10×10^-22) and CO25 (OR= 15.11, p-value= 2.83×10^-144). Similarly, we observed a consistent enrichment of S-phase cycling NPCs (ie. ‘PgS’) amongst up-regulated genes identified in ARTHS at CO15 (OR= 6.80, p-value=9.12×10^-38) and CO25 (OR= 13.70, p-value= 1.38×10^-160). (I) Protein-protein interaction network of key transcription factors driving (i.e. activating E2Fs, pluripotency) the expression of cell-fate modulating cell-cycle genes; these key transcription factors and their target genes are specifically upregulated in ARTHS during CO differentiation; network edges indicate both functional and physical associations, while the thickness of edges corresponding to strength of associations.

Notably, we also discovered that the gene expression of “activating E2Fs”, *E2F1, E2F2,* and *E2F3* (**Fig. 4B-D**), was significantly upregulated across multiple CO stages, but not the IPSC stage in ARTHS. In particular *E2F3* was overexpressed in ARTHS COs at both CO stages (**Fig. 4D**) and DNA motifs corresponding to this transcription factor were enriched amongst the 980 genes which were sigDE at both CO stages (**Fig. 4A**). *E2F3* is a regulator of NPC properties (e.g. patterning, differentiation, neurogenesis, cell fate) during ND through modulation of cell cycle dynamics ^97,98,103–105^. Irrespective of *KAT6A* mutation status, we also noted that *E2F3* expression decreased as IPSCs differentiated into COs. Strikingly, the *E2F3* expression pattern we observed throughout CO differentiation in both ARTHS and control samples, mirrors the *E2F3* expression pattern found *in vivo* within the developing mouse brain—where *E2F3 gene* expression is downregulated as neural differentiation proceeds^106^. To understand the biological relevance of the *E2F*- and cell cycle-transcriptional signatures in ARTHS COs, we determined the direction of change for a minimally-redundant lists of E2F targets (ID#M5925) and KEGG cell cycle genes (ID#M5925) obtained from the Human Molecular Signatures Database ^107–109^. We identified 128 up-regulated sigDEGs in the E2F target gene list (**Fig. 4E**) and 56 up-regulated sigDEGs in the KEGG cell cycle gene list (**Fig. 4F**). Interestingly, we only observed this significant regulation of E2F target genes and cell-cycle genes after the onset of neural differentiation in ARTHS COs (**Tables S8 and S10**). No differences in these E2F-related and Cell-Cycle-related gene sets were observed in ARTHS iPSCs (**Table S6**).

Since we observed significant enrichments of cell-cycle related genes amongst the genes which were sigDE only in COs (**Fig. 2**, **Fig. 4A-F**) we hypothesized that there is a significant difference in the expression of proliferative cell populations found in the developing human brain. We performed a cell type enrichment analysis on genes that were sigDE in ARTHS using 16 cell-type specific markers defined in a single-cell transcriptomic study of the human fetal brain ^42^. We identified several brain cell types that were enriched or depleted in ARTHS (**Table S16**). Concordant with our previous observations (**Fig. 2**, **Fig. 4A-F**), we found sigDEGs identified in ARTHS COs were significantly enriched for markers of cycling NPCs (**Fig. 4G-H**). Specifically, as shown in **Fig. 4G-H**, we identified a consistent enrichment of G2/M phase cycling NPCs (ie. ‘PgG2M’) amongst up-regulated genes identified in ARTHS at CO15 (Fisher’s Exact test: OR=5.47, p-value=2.10×10^-22) and CO25 (OR= 15.11, p-value= 2.83×10^-144). Similarly, we observed a consistent enrichment of S-phase cycling NPCs (ie. ‘PgS’) amongst up-regulated genes identified in ARTHS at CO15 (OR= 6.80, p-value=9.12×10^-38) and CO25 (OR= 13.70, p-value= 1.38×10^-160).

In order to confirm that *KAT6A* is expressed in the cell types that displayed enrichment in ARTHS, we plotted this gene’s expression within single cells of the developing human neocortex during mid-gestation using CoDex, a transcriptomic atlas developed by the authors that defines cell type markers used in the enrichment analysis^42^. This analysis showed that *KAT6A* and *KAT6B* are both broadly expressed across all 16 cell types identified in the previous study (**Fig. S16A**). Then to confirm the cell-cycle-related enrichment we observed in ARTHS sigDEGs, we plotted the normalized gene expression of cycling NPCs markers across all 3 stages of CO differentiation using our RNA-seq data and found that ARTHS COs display higher expression of ‘PgG2M’ and ‘PgS’ markers compared to unaffected controls (**Fig. S16B**).

Next, we investigated the roles of all the non-*E2F* DNA motifs that are significantly overrepresented amongst the promoters of the 980 sigDEGs which were persistently dysregulated in ARTHS COs (**Fig. 4A**) and found the non-*E2F* motifs correspond to TFs that play central roles in modulating neural differentiation and cell specification during ND and/or have been implicated in increasing susceptibility of ASD ^110–122^. One of the most noteworthy non-*E2F* TFs from this list is *ZFP57* (**Fig. 4A**) because: (1) this gene compromises ND pathways in ASD by contributing to the epigenetic load of this NDD ^123^ and (2) we observed motifs corresponding to this TF were also significantly overrepresented amongst the 235 genes which were consistently sigDE across all 3 stages of CO differentiation in ARTHS (**Fig. 3B**, **Tables S12 and S17**). Additionally, we identified a significant overrepresentation of *PBX1* motifs amongst the genes that are transcriptionally dysregulated in ARTHS Cos (**Fig. 4A**). *PBX1* is known to be regulated by a master RBP called *PTBP1* during neuronal differentiation ^122^ and this prompted us to investigate RBPs and their targets.

RBPs regulate RNA splicing in the developing brain to modulate cellular processes like cell identity transition ^46–52,55,124^. Our results showed an enrichment of RNA-processing genes (**Fig. 2E-F**) and *PBX1* motifs amongst genes that are sigDE in ARTHS COs (**Fig. 4A**)—so we hypothesized that aberrant transcription in ARTHS COs was driven by RBPs. To this end, we compared the lists of sigDEGs identified in ARTHS by srRNA-seq (**Tables S6, S8, S10**) to publicly-available RBP-related datasets (**Table S18**) and identified 50 RBPs that were sigDE in ARTHS across the 3 stages of CO differentiation (**Fig. S17A**, **Table S19**), with 34 sigDE RBPs displaying enrichment of their target genes in the ARTHS sigDE genes (**Fig. S17B**, **Table S20**). Interestingly, the total number of RPBs, and RBPs with target enrichment, increases in the up-regulated ARTHS genes as differentiation proceeds (**Fig. S17A-B**). The most striking finding in this RBP-target enrichment analysis was the consistent and significant overrepresentation of *PTBP1*-target genes up-regulated in ARTHS during all 3 timepoints (**Fig. S17C**, **Table S21**): IPSC stage (OR = 1.874, p-value= 3.45×10^-15), CO15 stage (OR = 1.430, p-value= 4.45×10^-05), and CO25 stage (OR = 1.712, p-value= 1.20×10^-24).

### Pluripotency markers have prolonged expression in ARTHS COs during neural differentiation

Lastly, we looked at the expression of classical NPC and pluripotency markers in ARTHS across all stages of CO differentiation. From this RNA-seq differential expression analysis, we found that the expression of NPC markers (i.e. PAX6, NESTIN) increased as the time of differentiation increased for all samples, and we detected no significant difference in expression for these NPC markers between the ARTHS and control samples (**Fig. S18**). The gene expression of 6 core pluripotency markers decreased for all samples as the time of differentiation increased (**Fig. S19**). However, we found that the gene expression of these 6 core pluripotency markers remained significantly elevated in ARTHS COs compared to control COs (**Fig. S19**). Specifically, at day 15 of CO differentiation, we found that ARTHS COs expressed significantly higher amounts of several core pluripotency genes compared to unaffected control: *NANOG* (Log_2_FC of ARTHS/control = 5.50, P_adj_ = 1.93×10^-4^), *OCT3/4* (Log_2_FC= 3.68, P_adj_= 6.40×10^-3^), *MYC* (Log_2_FC= 1.10, P_adj_ = 2.98×10^-2^), *SALL1* (Log_2_FC= 0.778, P_adj_= 1.75×10^-7^), *SALL4* (Log_2_FC = 0.799, P_adj_= 2.74×10^-13^), *HDAC1* (Log_2_FC = 0.401, P_adj_=2.34×10^-2^), *HDAC2* (Log_2_FC = 0.286, P_adj_= 3.91×10^-2^), and *TRIM28* (Log_2_FC = 0.338, P_adj_= 1.87×10^-2^). The expression of most of these pluripotency genes remained significantly elevated in ARTHS COs at day 25 of differentiation.

The gene expression of activating E2Fs (**Fig. 4B-D**) is known to drive gene expression of pluripotency (**Fig. S19**) and cell cycle-related (**Fig. 4E-F**) genes. Thus, in **Figure 4I**, we created a protein-protein interaction (PPI) network between all the activating E2Fs (**Fig. 4B-D**) and pluripotency genes that were specifically upregulated in ARTHS COs (**Tables S6, S8, S10**) to illuminate the interconnection between these two groups of TFs which precisely regulate cell identity transition from a pluripotency state to a differentiated state during neural differentiation—in tandem with RBPs like *PTBP1* (**Fig. S17**). Taken together, this data demonstrates that during the first 25 days of CO differentiation, when compared to controls, ARTHS COs exhibit a significant delay in repressing the expression of genes related to: pluripotency, cell cycle dynamics, activating *E2F*s, *E2F*-targets, *PTBP1*-targets, and markers of cycling human NPCs.

### Pathogenic KAT6A mutations disrupt transcription of Autism Spectrum Disorder (ASD) Risk Genes

ARTHS patients are often co-diagnosed with ASD, and in **Fig. 4A**, amongst the promoters of all the genes that are persistently dysregulated in ARTHS COs, we identified a significant overrepresentation of a motif corresponding to a gene implicated in the epigenetic load of ASD (i.e. ZFP57)^123^. We asked whether there was substantial overlap between the ARTHS sigDE genes and genes associated with ASD susceptibility (a.k.a ASD risk genes).

To illuminate the shared factors underlying ARTHS and ASD, we compared our ARTHS RNA-seq differential expression results to an evidence-based list of Autism spectrum disorder risk (ASDR) genes curated by the Simons Foundation Autism Research Initiative ^125,126^. Specifically, in these enrichment tests we stratified the list of SFARI ASDR genes based on their “Gene Scoring” value, considering either: all ASDR genes (1+2+3+S), high confidence ASDR genes (1+S) (**Fig. 5A**) or a subset of common (non-syndromic) (1+2) ASDR genes (**Fig. 5B**). Of the 1,066 ASDR genes expressed in our RNA-seq data (**Table S22**), 284 genes were sigDE in ARTHS in at least 1 out of 3 stages assessed via RNA-seq, which we hereafter refer to as “any-stage ASDR-sigDE” genes (**Fig. S20A**, **Table S23**). These 284 “any-stage ASDR-sigDE” were inclusive of both significantly up- and down- regulated genes and our enrichment analysis revealed these genes are significantly overrepresented (Fisher’s Exact: Odds Ratio (OR)= 1.86, p-value= 7.76×10^-17^, **Fig. 5A**) amongst all the 6,071 sigDE genes perturbed by pathogenic *KAT6A* mutations during CO differentiation (**Fig. 2**). In total, 4.67% of all sigDE genes that are either up- or down- regulated in ARTHS are significantly overrepresented in all the SFARI ASDR genes (284 of 6,071; OR= 1.86, p-value <0.05, **Fig. 5A**). Even when restricting to the high confidence SFARI ASDR genes, the enrichment remained significant (OR=1.85, p-value=9.19×10^-17^, **Fig. 5A**). Although the total number of sigDE ASDR genes varied across the 3 stages, we found ASDR genes are consistently overrepresented amongst genes that were sigDE in ARTHS at the start and end of CO differentiation (**Fig. 5A**).

**Fig 5:**
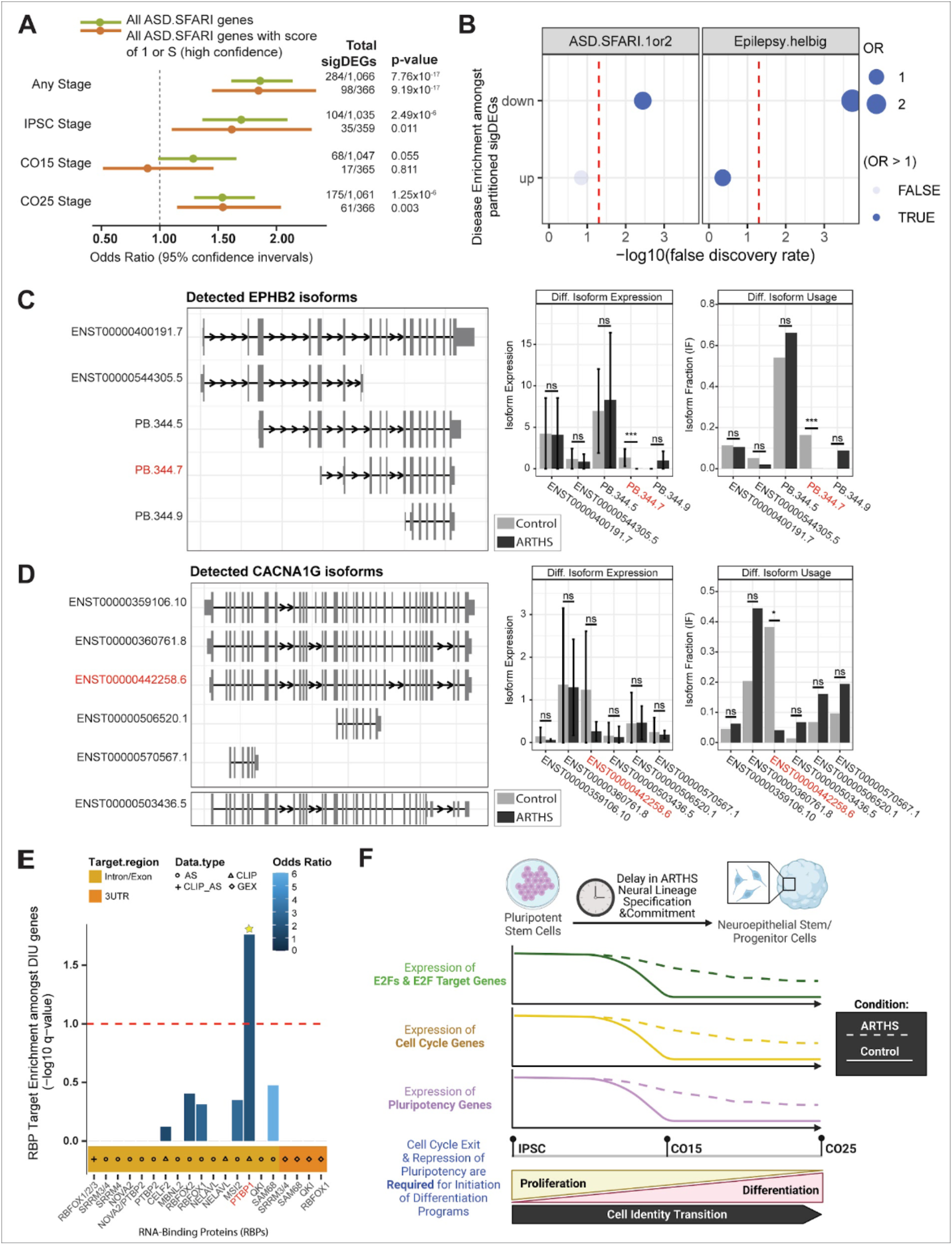
Pervasive transcriptional dysregulation of Autism risk genes discovered in ARTHS. (A) All the genes that were sigDE in ARTHS at any stage (Fisher’s Exact Test: Odds Ratio (OR) =1.86, p= 7.76×10^-17), the IPSC stage (OR=1.70, p-value=2.49×10^-6) and the CO25 stage (OR=1.54, p-value= 1.25×10^-6) were significantly overrepresented (p-value<0.05) amongst the SFARI ASD gene list when considering all genes (green) and high confidence genes (SFARI scoring of 1 or S). (B) All the genes that are sigDE in ARTHS in the RNA-seq data were partitioning into up-regulated (up) or down-regulated (down) based on the direction of L2FC before performing rare variant enrichment on partitioned sigDEGs. Amongst the down-regulated sigDEGs identified in ARTHS, there was a significant enrichment of ‘ASD.SFARI.1or2’ genes (Fisher’s Exact Test: OR = 1.49, FDR=3.60×10^-04). Similarly, amongst the down-regulated sigDEGs identified in ARTHS, there was a significant enrichment of ‘Epilepsy.helbig’ genes (OR = 2.73, FDR=9.32×10-06). No significant enrichment was detected amongst the up-regulated sigDEGs in ARTHS across either the ‘ASD.SFARI.1or2’ genes (OR = 0.839, FDR= 7.52×10^-02) or ‘Epilepsy.helbig’ genes (OR = 1.23, FDR=3.51×10^-01). Next in our isoform analysis, we identified two epilepsy-related ASD risk genes, (C) EPHB2 and (D) CACNA1G, that both show significantly reduced usage of one specific RNA isoform in ARTHS-patient biosamples relative to controls (P_adj_ < 0.05, affected isoforms are in red). (C) EPHB2 exhibits both DIU and DIE in ARTHS as indicated by significantly reduced usage and expression of the isoform “PB.344.7” (dIF = −0.164 with a q-value of 1.62×10^-19, isoform L2FC =-7.08 with a q-value of 3.96×10^-08). (D) CACNA1G exhibits DIU in ARTHS as indicated by significantly reduced usage of the isoform “ENST00000442258.6” (dIF = −0.342 with a q-value of 1.29×10^-03); transcripts starting with the prefix “PB” are novel RNA isoforms identified through PacBio’s Iso-Seq platform, while those starting with “ENST” are known RNA isoforms present in the GENCODE reference annotation and Y-axis of ‘isoform expression’ is the L2FC in the DIE analysis. (E) The enrichment of brain-specific RBP targets was evaluated amongst the top DIU hits. This analysis revealed that *PTBP1-*target genes were enriched amongst the genes that displayed significant DIU in ARTHS via lrRNA-seq (Fisher’s Exact Test: OR = 1.93, p-value=1.73×10^-02). (F) Proposed model by which pathogenic *KAT6A* mutations present in ARTHS-patient COs drive aberrant transcription of proliferation-related gene networks known to modulate cell fate acquisition in neural lineages. Importantly, aberrant cell cycle/proliferation dynamics are a common disease-associated feature found in ASD and they are thought to contribute to observed clinical phenotypes.

To holistically classify the function(s) of these 284 any-stage ASDR-sigDE genes (**Table S23**), we performed a GO analysis on this gene list for BP, CC, and MF terms (**Table S24**). In the first GO analysis, we found that these any-stage sigDE ASDR genes are significantly enriched (P_adj_ < 0.05) for 334 BP GO terms that are predominantly related to: brain development, and cerebrum function (e.g. learning, memory, cognition) and neuron physiology; the top 7 BP GO terms relate to membrane potential, synaptic signaling, and synapse organization (**Figs. S21-S22**). In the second GO analysis, we found the 284 any-stage ASDR-sigDE genes are significantly enriched (P_adj_ < 0.05) for 132 CC GO terms that mainly related to: synapse physiology, biomolecule complexes (e.g. ion channel complex), and cell membrane dynamics (**Figs. S23-S24**). In the third GO analysis, we found the 284 any-stage ASDR-sigDE genes are significantly enriched (P_adj_ < 0.05) for 72 MF GO terms related to ion channel activity, neurotransmitter receptor activity, and biomolecule binding (**Figs. S25-S26**). Other significantly overrepresented GO terms include those related to (**Table S24**): histones, DNA or chromatin or chromosomes, and the cardiovascular system (e.g. blood, heart, cardiac). Collectively, this shows that pathogenic mutations in *KAT6A* that cause ARTHS cause transcriptional dysregulation at SFARI ASD risk loci compared to controls. These dysregulated genes directly relate to several fundamental aspects of neurobiology like cognition and ion channel activity (**Table S23-S24**, **Figs. S21-S26**).

Ion channel dysregulation is known to underlie Epilepsy ^127^, an NDD that often co-occurs in both ARTHS^17^ patients and ASD ^128,129^ patients. Therefore, we sought to dissect the directionality of gene expression because in the previous enrichments we considered all sigDE genes (i.e. both up- and down-regulated) in ARTHS at any-stage of CO differentiation (**Fig. 5A**, **Table S23-S24**, **Figs. S21-S26**). To this end, we tested the enrichment of high-confidence and common ASDR genes and Epilepsy-related genes in sigDEGs whose expression is down-regulated (i.e. decreased L2FC) or up-regulated (i.e. increased L2FC) in ARTHS. This enrichment analysis revealed there is a significant overrepresentation of both high confidence and common ASDR genes and Epilepsy-related genes in transcriptionally down-regulated genes in ARTHS (FDR<0.05, **Fig. 5B**).

Since phenotypic overlap occurs between unrelated NDDs^128–132^, we next sought to define how high confidence ASDR genes (SFARI score = 1+S) that are significantly transcriptionally dysregulated in ARTHS across any stage of CO differentiation (n= 98 genes) relate to human phenotypes and how these genes physically interact with each other in protein-protein networks. We constructed a protein-protein interaction (PPI) network with all 98 high-confidence ASDR-sigDE genes using a highly-cited database called STRING ^133,134^. From this STRING analysis we found that these 98 genes were enriched for PPIs (*p-value* < 1.0×10^-16^) and 87.8% (86/98) had at least one PPI with confidence ≥ 0.4.

Moreover, from the Human Phenotype Ontology (HPO) enrichment analysis performed by STRING, we determined this gene network is significantly enriched for 692 HPO terms (False Discovery Rate (FDR) < 0.05; **Table S25**). Since this gene network was identified by overlapping genes that are sigDE in ARTHS with ASDR genes, we expectedly found a significant overrepresentation of these 98 genes amongst ASD-related HPO terms including: “Restrictive behavior” (HP:0000723), “Impaired social interactions” (HP:0000735), and “Poor eye contact” (HP:0000817). We unexpectedly discover that a large proportion of these 98 high-confidence ASDR-sigDE genes (35.7% to 67.3%) were significantly overrepresented in at least 1 of 13 HPO terms related to clinical manifestations in ARTHS (**Fig. S20**).

To elucidate the phenotypic interconnection between these 98 ASD risk genes that are transcriptionally dysregulated in ARTHS, we then calculated the odds of this high-confidence ASDR-sigDE gene network being present in gene networks of the 692 HPO terms first identified by STRING relative to all the genes present in the STRING database (**Table S25**). From this odds ratio analysis, we observed a significant overrepresentation of these 98 ASD risk genes (FDR < 0.05) amongst the 13 ARTHS-related HPO terms (**Fig. S20B**). We then inquired if any of these 98 ASD risk genes had been previously associated with >75% of the ARTHS-related HPO terms (i.e. 10 of 13 HPO terms shown in **Fig. S20B**) to evaluate their relationship to ARTHS and *KAT6A*. From this inquiry, we found 36/98 of high-confidence ASDR-sigDE genes (36.7%) are associated with at least 10 ARTHS-related HPO terms—and 30/36 of these genes form a robust PPI network (**Fig. S20C**), indicating these genes function within convergent biological pathways. Given their association with >75% of the ARTHS-related HPO terms, we next looked to see if these genes had been identified by OMIM as genetic causes of mendelian disorders. To our surprise, 35/36 (97.2%) of high-confidence ASDR-sigDE genes that exhibit a high degree of phenotypic overlap with ARTHS-related HPO terms also cause at least one monogenic developmental disorder (**Table S26**). Taken together, our findings provide further experimental evidence that there must be core biological pathways that converge to drive clinical phenotypes commonly manifested in genetic developmental disorders.

### Long-read IsoSeq analysis identify isoform-specific dysregulation in ARTHS samples

The enrichment of RNA-processing genes (**Fig. 2E-F**), RBP-target genes (**Fig. S17**), and ASDR genes (**Fig. 5A-B**) within the sigDEG identified in ARTHS biosamples led us to ask whether any of these categories showed isoform-specific transcriptional changes in ARTHS. These subtle changes in the transcriptome are difficult to detect by short-read RNA-seq alone as the read length only covers a fraction of a RNA transcripts. We performed long-read RNA-seq on ARTHS and control samples at the iPSC and CO25 differentiation using PacBio IsoSeq technology to sequence full-length transcripts for detection of differential isoform usage (DIU) and differential isoform expression (DIE) from the data using an R package called IsoformSwitchAnalyzeR ^135^. We identified 174 DIU genes, with or without co-occurrence of DIE, as indicated by differential isoform fractions (dIFs, q-value< 0.05, **Table S27**).

Of these 174 DIU genes, we found 7 were ASDR genes that exhibited significant DIU of at least 1 isoform in ARTHS (**Fig. 5C-D, Fig. S27A-E**): *EPHB2* (SFARI score = 2), *CACNA1G* (SFARI score =2), *SAMD11* (SFARI score = 2), *MACROD2* (SFARI score = 2), *CPZ* (SFARI score = 2), *USP30* (SFARI score = 3), and *PRKN* (SFARI score = 2). Epilepsy-related genes were significantly down-regulated in ARTHS CO (Fig. 5B), as shown in **Figure 5C-D**, and we identified two genes: *EPHB2* (dIF = −0.164, q-value of 1.62×10^-19) and *CACNA1G* (dIF = −0.342, q-value of 1.29×10^-03), that exhibited significantly reduced isoform usage in ARTHS. *EPHB2* encodes a developmentally-regulated Ephrin receptor that regulates NPC proliferation, neuronal differentiation, and neural tube closure ^136–139^ and mutations are associated with epilepsy ^140–142^. *CACNA1G* encodes a voltage-sensitive calcium channel that is an epilepsy-related gene ^143–149^ that regulates *in vitro* neural activity in both neurons and COs ^150,151^.

Next, since we previously observed a consistent significant overrepresentation of *PTBP1*-target genes amongst up-regulated sigDEGs identified in ARTHS during all 3 stages of CO differentiation (**Fig. S17, Table S21**)—we asked whether isoforms exhibiting DIU in ARTHS would also be enriched for RBP-target genes. To this end, we compared the list of top DIU genes identified in ARTHS by RNA-seq (**Table S27**) to the same lists of RBP-target genes (**Table S18**). From this comparison, we identified 105 RBPs that exhibited enrichment of their target genes amongst the isoforms that displayed differential usage in ARTHS via long-readRNA-seq (**Table S28**). In agreement with our findings in **Figure 5E**, *PTBP1*-target genes were also enriched amongst the genes that displayed significant DIU in ARTHS via Isoseq (Fisher’s Exact Test: OR = 1.93, p-value=1.73×10^-02).

We propose a model by which pathogenic *KAT6A* mutations in ARTHS COs drive aberrant transcription of proliferation-related gene networks to modulate neural differentiation of NPC lineages during *in vitro* brain development. Importantly, in ASD NPCs, aberrant proliferation and altered cell cycle dynamics are thought to contribute to observed clinical phenotypes associated with ASD ^152–155^. Thus, the mechanisms underlying this proposed pathogenic proliferation-differentiation model in ARTHS COs (**Fig. 5F**) likely involve interactions between: *KAT6A*, the cell cycle, alternative RNA splicing, pluripotency, activating E2Fs, RBPs like *PTBP1*, and ASD risk genes.

## Discussion

The gene regulatory effects of *KAT6A* mutations have never been described in human brain-like tissue models created from ARTHS iPSCs. To address this gap in the scientific literature, we performed a time-course *in vitro* model system of early brain development (Fig. 1) from iPSCs derived from 2 ARTHS patients and age- and sex-matched iPSCs from unaffected controls. We used both short-read RNA-seq and long-read IsoSeq analysis to comprehensively profile the transcriptomic perturbations caused by truncating *KAT6A* mutations at 3 stages along CO differentiation (Figs. 2-5). This approach enabled us to identify aberrant molecular signatures that underlie the neurodevelopmental phenotype observed in ARTHS. For the first time, we define how pathogenic *KAT6A* mutations cause to widespread transcriptional dysregulation of: (1) core pluripotency genes (), (2) key biological programs and pathways required in normal embryonic development (Figs. 2-4), and (3) a network of ASD risk genes that modulate fundamental aspects of neurobiology like cognition and synaptic plasticity (Figure 5). Long-read Iso-Seq data from iPSC and CO differentiation support that *KAT6A* mutations are associated with altered global expression and isoform usage in autism risk genes.

This study aimed to uncover genes and gene networks whose transcriptional state was significantly perturbed by the presence of pathogenic mutations in *KAT6A* throughout the earliest stages of brain development. Understanding the drivers underlying the neuro-related clinical phenotypes in ARTHS and other monogenic NDDs will enable development of molecular biomarkers, therapeutic drug targets, and predict off-target effects.

### Haploinsufficiency of KAT6A in ARTHS iPSCs and ARTHS COs

A common mechanism underlying the phenotypes associated with autosomal dominant disorders is haploinsufficiency, where reduced expression or activity of the affected gene is insufficient for maintaining a normal phenotype ^157^. In the DECIPHER database ^158^ classifies *KAT6A* as a haploinsufficiency gene which means it likely causes pathogenicity upon gene-dosage imbalances. *KAT6A* haploinsufficiency has been proposed as a cause of clinical phenotypes in ARTHS by others who have found ARTHS patients with heterozygous deletion of the entire KAT6A gene display the same ARTHS phenotypes as patients with other types of heterozygous *KAT6A* mutations ^2,159^. In agreement with this previous literature in humans, our data suggests that the gene regulatory effects (i.e. molecular phenotypes) in ARTHS are caused by haploinsufficiency of KAT6A since, relative to controls, ARTHS samples harboring truncating *KAT6A* mutations exhibit significantly reduced KAT6A expression at the protein-level (Fig. 1C) and RNA-level (Fig. 3A). The occurrence of KAT6A haploinsufficiency in humans is consistent with findings in mice, where the mouse KAT6A gene functions in a dosage-dependent manner and reduced KAT6A expression (i.e. haploinsufficiency) causes abnormal phenotypes across multiple tissues ^86,88,160^. Our finding are also supported literature which shows developmental disorders can be caused by haploinsufficiency in genes encoding transcriptional regulators, like KAT6A, through perturbations to biological programs that are also transcriptionally dysregulated in ARTHS in our data (specification, differentiation, cell fate)^161^. Our data adds to the growing literature on KAT6A haploinsufficiency in both human and non-human model organisms.

### Delayed repression of key cell-cycle regulation genes causes impaired neural specification in early neural development

One of the core findings in this study is that the expected repression of pluripotency gene expression during differentiation is significantly delayed in ARTHS COs relative to unaffected control CO. The significant elevation of gene expression in key pluripotency genes in ARTHS samples during CO differentiation was unexpected given that, prior to differentiation, ARTHS and control iPSC lines expressed similar levels of these pluripotency genes at the RNA and protein levels (**Fig. 1C**). When differentiation is induced in pluripotent stem cells, it is typically thought that the gene expression of many pluripotency genes gradually decreases until they are suppressed to permit the upregulation of genes critical for differentiation ^162^. However, this view is not entirely accurate since the expression of pluripotency genes, like those found to be elevated in our ARTHS COs, functions as lineage-specific blockages during differentiation to permit the generation of a specific lineage over other possible cell fates ^163^. The latter may explain why we see a significant enrichment of gene modules associated with the specification, differentiation, and development of various non-neural embryonic tissues amongst all the genes that are most transcriptionally dysregulated in ARTHS COs. Although the exact mechanisms by which the human *KAT6A* gene regulates the expression of pluripotency genes were not identified here, our findings are the first to indicate that the human *KAT6A* gene plays a previously unrecognized role in modulating the expression of pluripotency genes to potentially influence the lineage choices of ARTHS iPSCs.

### Pathogenic KAT6A mutations cause widespread transcriptomic perturbations

Given KAT6A’s role in controlling the epigenome, we expect to have widespread transcriptional perturbations due to *KAT6A* mutations across all 3 stages of CO differentiation and our experiments identified a total of 6071 genes whose transcriptional state is significantly dysregulated in ARTHS compared to controls. One challenge in the use of cerebral organoids is that by the nature of a more physiological model with cellular heterogeneity, it becomes increasingly challenging to disentangle effects of cell-type composition from direct cell-intrinsic effects of KAT6A mutations. We were able to identify clear lineage-independent effects of pathogenic *KAT6A* mutations where we noted 235 genes are consistently sigDE in ARTHS across all 3 stages assessed by srRNA-seq (**Fig. 3B**). These lineage-independent transcriptional dysregulation observed in cells harboring pathogenic KAT6A mutations support the notion that KAT6A modulates gene regulatory effects that transcend cellular context.

Here we highlight and discuss the neurological implications of two genes, *CTSF* and *ZNF229*, that were identified as being significantly overexpressed in ARTHS at all 3 stages of CO differentiation compared to unaffected controls (**Fig. 3**). In ARTHS patients, the gene expression of *CTSF* is at least 4 times higher than unaffected controls (Log_2_FC ranging = 2.11-3.79, P_adj_ range = 3.44×10^-04^ - 5.07×10^-38^). While the gene expression of ZNF229 is at least 124 times higher in ARTHS patients compared to controls (Log_2_FC range = 6.96-7.56, P_adj_ range = 2.78×10^-59^-5.07×10^-138^)—making ZNF229 one of the most transcriptionally dysregulated gene in our RNA-seq data. Strikingly, both CTSF and ZNF229 have been previously linked to different aspects of neurobiology or embryonic development. *CTSF* encodes a cysteine protease that: (1) is normally ubiquitously expressed at high levels in the human brain during prenatal development into young adulthood, (2) is required for protein homeostasis in the brain, and (3) is known to cause an autosomal recessive mendelian neurodegenerative disorder called Kufs-type neuronal ceroid lipofuscinosis-13 (CLN13, OMIM#615362) which is characterized by progressive cognitive decline and motor dysfunction ^164,165^. Although this is beyond the scope of this study, we speculate that perhaps gene-dosage imbalances in *CTSF* may explain the impaired cognitive and motor function observed in ARTHS patients given that these phenotypes are also observed in CLN13. Next, *ZNF229* may also be an important contributor to ARTHS pathogenesis given that this gene encodes a hominoid-specific zinc finger protein (ZFP) which has been recently implicated in controlling the activation of transcriptional *cis* regulatory elements during early embryogenesis ^166^. Importantly, *ZNF229* has also been reported to be one of the most highly expressed genes specific to human neural crest lineages from *in vitro* ectodermal differentiation experiments^26^. The lineage-independent effects of pathogenic KAT6A mutations identified here in the context of ARTHS warrant further mechanistic studies to determine if any of these genes can be targeted to develop potential cross-tissue therapies for this NDD.

### ARTHS COs display aberrant transcription of genes known to govern cell fate during neurodevelopment

The significant transcriptional dysregulation observed in ARTHS COs is unlikely due to differences *in vitro* brain size given we did not observe a difference in the size between ARTHS and control COs at day 25 of differentiation. The 980 genes that are persistently sigDE in ARTHS COs across both stages of CO differentiation are significantly enriched in genes that affect cell fate in the developing brain by either modulating differentiation via cell cycle dynamics (i.e. E2F-related genes) (Fig. 4) or influencing of neural circuitry formation via adhesion molecules (i.e. PCDH genes). Our findings are supported by literature that identified aberrant molecular signatures in E2F-related genes ^1^ and PCDH genes ^167^ in ARTHS primary fibroblasts—again suggesting that some of the gene regulatory effects of pathogenic KAT6A mutations in ARTHS may transcend cellular context.

Taken together with our previous findings that ARTHS COs: (1) fail to downregulate the expression of key pluripotency genes and (2) display significant transcriptional dysregulation of gene modules associated with the specification, differentiation, and development of non-neural embryonic tissues ()—this consistent transcriptional dysregulation of E2F-related genes and PCDH genes observed in ARTHS COs (Fig. 4) demonstrate pathogenic KAT6A mutations impede proper *in vitro* brain development through multiple biological perturbations. Interestingly, in a different Mendelian developmental syndrome called Seckel syndrome (OMIM#210600), which is caused by germline mutations in the epigene ATR, researchers also found that patient COs exhibited differentiation defects due to dysfunctional cell cycle dynamics ^168^—suggesting that dysfunctional cell cycle dynamics may commonly affect neural differentiation in developmental syndromes like ARTHS.

### Effect of pathogenic KAT6A mutations on transcription of Autism Spectrum Disorder (ASD) Risk Genes

Even before the first association of *de novo KAT6A* mutations with the constellation of clinical features known as ARTHS, *KAT6A* was identified as a genetic risk factor for ASD^169^. In this study, we discovered significant transcriptional dysregulation of ASD risk genes in ARTHS patient samples across multiple stages of CO differentiation. The ASD risk genes that are significantly differentially expressed in ARTHS patients regulate fundamental aspects of early brain cell specification, setting the stage for broad disruption of normal neurodevelopment. This is the first study to explore the time-course transcriptional dysregulation in ARTHS patient-derived stem cell models. The enrichment of ASD risk genes within the differentially expressed gene set supports the presence of early, convergent molecular pathways driving neurodevelopmental delay. We suspect that the early effects of *KAT6A* in neural specification might be common to other organ systems affected by KAT6A. Although NDDs have divergent genetic etiologies, these genes all disrupt a common sequence of events: cell proliferation, cycle, migration, or differentiation, resulting in a spectrum of clinical neurodevelopmental phenotypes that include ASD. We do not believe it is surprising that often epigenes can be associated with multiple neuropsychiatric traits, highlighting the redundant signaling pathways in neurodevelopment. We show that decreased expression of chromatin-related ASD risk genes (e.g. KAT6A haploinsufficiency), dysregulates molecular signatures that converge on pathways which modulate cell-cycle dynamics and the temporal timing of neural differentiation ^170^ and highlight potential targets to repairing neuroconnectivity in childhood and beyond.

## Materials and Methods

### Institutional Approvals and human iPSC Generation

This study was approved by the Institutional Review Board and the Embryonic Stem Cell Research Oversight (ESCRO) committees at UCLA. All biological samples were collected after informed consent (IRB#11-001087). Whole blood or dermal fibroblasts were submitted to the Cedars-Sinai Regenerative Medicine Institute’s Induced Pluripotent Stem Cell Core Production Facility for conversion into iPSCs using episomal reprogramming with the following factors: OCT3/4, SOX2, KLF4, L-MYC, SHP53, and LIN28. As a part of their reprogramming quality control, Cedar-Sinai performed G-band karyotyping, pluripotency immunocytochemistry, alkaline phosphatase staining, and IDEXX cell line authentication. Pluripotency status of all iPSC lines was re-assessed in-house prior to performing differentiation experiments.

### Maintenance of human iPSC lines

All iPSC lines used in this study were grown under chemically-defined and feeder-free conditions in 37□ incubators supplemented with 5% CO_2_ at atmospheric concentrations of O_2_. All iPSC lines were maintained on Nunc 6-well culture plates (ThermoScientific #140675) coated with recombinant human vitronectin protein at a final concentration of 0.50 ug/cm^2^ (Gibco #A14700) in complete Essential 8 Medium (complete E8; Gibco #A1517001). A complete media change was performed daily to prevent spontaneous differentiation. Plates were coated as follows: first stock vitronectin (0.5 mg/mL) was diluted 1/100 in DPBS without calcium or magnesium (Gibco #14190144), then diluted vitronectin was immediately dispensed into plates, followed by either overnight or 2 hour incubation at 37□ prior to use. Stock vitronectin (0.5 mg/mL) was aseptically aliquoted into 1.5 mL polypropylene tubes and stored at −80□.

All iPSC lines were passaged as clumps or single cells upon reaching 80-85% confluency using Versene (0.02% EDTA solution, Lonza #17-711E) or StemPro Accutase (Gibco #A1110501). To ensure optimal cell survival, for the first 12-18 hours after passaging, iPSC lines were fed complete E8 medium supplemented with Y-27632 dihydrochloride at final concentration of 10uM (Tocris Bioscience #1254). To create a 1000X stock, Y-27632 powder was reconstituted in sterile nuclease-free water to a stock concentration of 10mM before being filtered (MilliporeSigma #SE1M179M6) and aliquoted into sterile 1.5 mL polypropylene tubes which were then stored at −80C. All cultures in this study were routinely tested for mycoplasma contamination per manufacturer instructions (Lonza #LT07-318).

### KAT6A genotyping of human iPSC lines via Sanger sequencing

To confirm cell line identity, all iPSC lines used in this study were genotyped prior to cerebral organoid differentiation. For each iPSC line, 1 million cells were washed, pelleted and then snap-frozen on dry ice. Snap-frozen iPSC pellets were extracted using a DNeasy Blood & Tissue Kit per manufacturer instructions (Qiagen #69504) and eluted in sterile nuclease-free water (Invitrogen #10977023). The quality and quantity of extracted DNA was assayed using a NanoDrop Spectrophotometer (Thermo Scientific #701058112); all extracted DNA exhibited 260/280 values greater than 1.8 and 260/230 values greater than 2.0, indicating pure DNA extractions.

Genotyping primer pairs were designed to flank both known ARTHS patient mutations using NCBI’s genomic reference sequence for KAT6A (NG_042093.1 RefSeqGene). To genotype exon 7 of KAT6A, the following primers were used: 5’-TCTGACTCCTGGCCTG-3’ and 5’-CACCTCAGTATCATCTACATG-3’. To genotype exon 17 of KAT6A, the following primers were used: 5’-AGGAGGAGGAAGATGCAG-3’ and 5’-TCACTGAAGCCGCTGT-3’. These primers were purchased as single stranded DNA oligonucleotides from Integrated DNA Technologies (IDT) and were initially reconstituted to 100uM in 1X IDTE (IDT #11-01-02-02) before diluting further to create 10uM stock primers that were used for polymerase chain reaction (PCR). The primers used to genotype exon 7 and exon 17 of KAT6A were expected to produce amplicons of 711 and 760 base pairs in length, respectively.

To empirically identify which PCR conditions would yield a single amplicon of expected length, we first performed a series of gradient PCRs with control DNA using: the aforementioned primer pairs, OneTaq DNA polymerase master mix (New England Biolabs #M0489S), and a T100 Thermal Cycler (Biorad #1861096). For all extracted DNA and genotyping primer pairs, the following PCR conditions were used to generate a single KAT6A amplicons: (1) OneTaq mastermix was added to a final concentration of 1X, (2) DNA was added to a final concentration of 5 ng/uL, (3) forward and reverse primers were each added to a final concentration of 0.2uM, and (4) nuclease-free water was added to bring the PCRs up to the final desired volume. Also, for all extracted DNA and genotyping primer pairs, the following thermocycler conditions were used to generate a single KAT6A amplicons: an initial denaturing at 94C for 45 seconds, 35 cycles of 94C for 20 seconds + 50C for 30 seconds + 68C for 50 seconds, and a final extension at 68C for 5 minutes. To confirm the presence of a single amplicon of expected length prior to sequencing, the resulting PCR reactions were run on a 2% agarose/TAE gel (Thermo Scientific #B49), UltraPure Agarose (Invitrogen #16500500), and 1X SYBR Safe DNA Gel Stain (Invitrogen #S33102). The resulting gel was then visualized on a ChemiDoc Imaging System (Biorad #17001401) with a blue tray (Biorad #12003027).These single-amplicon PCRs were then purified using a QIAquick PCR Cleanup Kit (Qiagen #28506) before submitting to Genewiz (La Jolla, CA) for Sanger sequencing. Lastly, sanger sequencing results were aligned to the genomic sequence of KAT6A using SnapGene software (www.snapgene.com).

### Human iPSC differentiation into Cerebral Organoids

Human iPSC lines derived from ARTHS patients and unaffected controls were differentiated for 25 days into early cerebral organoids using a previously published protocol ^70^. To this end, we referenced the methods provided in the original publication in conjunction with a step-by-step guide posted on Protocol Exchange by the same authors ^171^. The optional pre-treatment of iPSCs with DMSO was not used in this study and we also did not include the optional XAV-939 compound during the first 5 days of cerebral organoid differentiation. To prevent mechanical shearing of iPSC-derived spheroids and cerebral organoids, we added the following procedures throughout this protocol: (1) we pretreated all plasticware that came into contact with cell aggregates with an anti-adhesion solution (Stem Cell Technology #07010) per manufacturer instructions and (2) we used wide-bore serologicals and wide-bore pipette tips to handle cell aggregates.

To create iPSC-derived spheroids for cerebral organoid differentiation, iPSCs were first grown on vitronectin-coated plates in complete E8 Medium as previously described in this paper. Once iPSCs reached ∼90% confluency, cultures were dissociated into single cells using accutase. Single-cell suspensions were then quantified in triplicate using an automated cell counter (Invitrogen #C10228) per manufacturer instructions. Next, to form consistent iPSC-derived spheroids, three million iPSCs were seeded into each well of an AggreWell-800 plate (Stem Cell Technology #34815), resulting in ten thousand cells per microwell. Seeding iPSCs into AggreWell-800 plates containing complete E8 medium supplemented 10uM Y-27632 marked day 0 of spheroid formation. Lastly, to ensure efficient generation of iPSC-derived spheroids, cells were cultured in AggreWell-800 plates for two days, with daily media changes, before harvesting for differentiation.

Subsequently, iPSC-derived spheroids were harvested into neural induction medium (NIM) and cultured in ultra-low attachment plates (Corning #3262), marking day 0 of differentiation. As described in the original protocol, cultures were fed fresh NIM from day 0 to day 5 of cerebral organoid differentiation. This NIM consisted of Essential 6 medium (Gibco #A1516401) supplemented with Dorsomorphin (Sigma Aldrich #P5499-5MG) and SB-431542 (Tocris Bioscience #161410) at a final concentration of 2.5 uM and 10 uM, respectively. From day 6 to day 25 of differentiation, cultures were grown in neural maturation medium (NMM) as described in the original protocol to promote cerebral organoid growth and maturation. This NMM was composed of Neurobasal-A Medium (Gibco #10888022) supplemented with the following reagents at the specified final concentrations: B-27 at 1X (Gibco #12587010), GlutaMAX at 1X (Gibco #35050061), recombinant human EGF at 20 ng/mL (R&D systems #236-EG-200), and recombinant human FGF2 at 20 ng/uL (R&D systems #233-FB-010).

### Histology of Cerebral Organoids

As previously mentioned, all plasticware used to handle the cerebral organoids was pretreated with an anti-adhesion solution (Stem Cell Technology #07010). All fixation steps described in this study were performed using 4% paraformaldehyde (PFA) that was prepared in-house as follows. To make 500mL of 4% PFA, first prepare 1X PBS (Fisher Scientific #BP3994) using Milli-Q water (Millipore #Z00Q0V0WW). Next, add 20 grams of PFA crystals (Sigma #P6148-500G) and 2 NaOH chips (Fisher Scientific #S318-500) to 300mL of 1X PBS in a glass beaker. To dissolve the PFA crystals, place the PFA solution on a hot plate and mix using a magnetic stir bar until the solution becomes clear; the temperature of the PFA solution should not exceed 60C. Allow the solution to cool to room temperature, then measure the starting pH of the PFA solution. Next, while stirring without heat, perform an acid-base titration on the PFA solution by carefully adding either HCl or NaOH dropwise as necessary until a pH of 7.4 is reached. Then, bring the PFA solution up to a final volume of 500mL by adding more 1X PBS. Lastly, filter the resulting 4% PFA solution with Whatman paper (Sigma #WHA1003185) before aliquoting and freezing at −20C.

At day 25 of cerebral organoid differentiation, biological specimens were individually collected over 37 um filters (Stem Cell Technology #27215) using a dissecting microscope and sterile P1000 pipette tips that were manually cut with sterile disposable scalpels (Fisher Scientific #08-927-5B). Samples were washed once with ice-cold DPBS (Gibco #14190144), followed by overnight fixation at 4C using ice-cold and freshly-thawed 4% PFA. The following day, cerebral organoids were washed 4 times in ice-cold DPBS at 4C to remove excess PFA. Fixed cerebral organoids were then dehydrated at 4C in ice-cold 70% ethanol (Fisher Scientific #BP2818500) overnight. To prevent sample loss during downstream steps, dehydrated and fixed cerebral organoids were embedded in histology gel (Epredia #HG4000012) per manufacturer instructions. Once the histology gel cooled and solidified, the gel-embedded cerebral organoids were then transferred into tissue-embedding cassettes and stored at 4C in ice-cold 70% ethanol. The following day, these cassettes were submitted to UCLA’s Translational Pathology Core Laboratory (TPCL) for paraffin embedding, sectioning, and mounting. Paraffin blocks were sectioned at a thickness of 5 um before mounting on glass slides. A subset of paraffin blocks were also processed by UCLA’s TPCL for Hematoxylin and Eosin (H&E) staining, obtaining an average of 18 H&E slides per cell line, with 3 to 4 sections per slide.

### QuPath Size Analysis of H&E-stained cerebral Organoids

All H&E slides were digitally scanned at 40x magnification by UCLA’s TPCL using the brightfield setting on an Aperio ScanScope AT system. All resulting brightfield images underwent quality control to ensure the whole H&E slide was scanned and in focus; this imaging data was returned to us as ‘.svs’ files. Next, to determine the average size of each cerebral organoid after 25 days of differentiation, we counted the number of cells present in each H&E section using QuPath software v0.2.3 ^71^. Specifically, QuPath’s pixel classifier was first used to detect and delineate individual organoids in each slide section. Then QuPath’s cell detection function was used to identify and count individual cells within each H&E section. Lastly, we evaluated whether there was a significant statistical difference between the average size of ARTHS COs and control COs using a standard t-test. The groovy script used in this QuPath analysis is provided in the supplementary material.

### Immunocytochemistry (ICC) and Immunohistochemistry (IHC) Microscopy

Undifferentiated iPSC lines were grown as previously described on vitronectin-coated glass coverslips or tissue culture plates. A 2-day immunocytochemistry (ICC) protocol was then performed on iPSCs by first washing cells once with ice-cold DPBS (Gibco #14190144), followed by fixation of cells via incubation with freshly-thawed 4% PFA for 15 to 20 minutes at room temperature. To remove excess PFA, cells were then washed once with ice-cold DPBS. Next, cells were permeabilized and blocked by incubating with 1X PBS containing 5% donkey serum (Jackson Immuno Research #017-000-121) and 0.1% triton×100 (Sigma #X100-1L) for 20 to 30 minutes at room temperature. All primary antibody solutions were then prepared in 1X PBS containing 5% donkey serum and allowed to incubate with cells overnight at 4C. Cells were co-stained with a rabbit anti-NANOG (Cell Signaling #4903S) and a mouse anti-OCT3/4 (Santa Cruz #SC5279) antibody using a 1/200 dilution for each. The following day, to remove excess antibodies, cells were washed at room temperature 3 times in 1X PBS with gentle agitation for 5 minutes for each wash. All secondary antibodies were diluted in 1X PBS containing 5% donkey serum and 0.001mg/mL Hoechst (Invitrogen #H3570) before allowing cells to incubate with said solution for 1 hour at room temperature. Cells were co-stained with a donkey anti-rabbit-AlexaFluor488 (Invitrogen #A32790) and donkey anti-mouse-AlexaFluor555 (Invitrogen #A32773) antibody using a dilution of 1/500 for each. Next, cells were washed in 1X PBS as previously described to remove excess antibodies. Lastly, cells were imaged at 40X on a fluorescent Zeiss Axio Vert-A1 microscope.

We performed immunohistochemistry (IHC) on unstained paraffin-embedded COs isolated on day 25 of differentiation using a modified protocol ^172^. The unstained paraffin-embedded tissue was fixed, embedded, sectioned, and mounted onto glass slides as previously described in the CO histology section. Briefly, on the first day, the slides were: (1) deparaffinized and rehydrated with a series of incubations in xylene, ethanol, and water, (2) permeabilized and blocked, and then (3) incubated overnight with primary antibodies at 4C. On the second day, the slides were: (4) washed in 1XPBS three times for 10 minutes each wash, (5) incubated for 2 hours with secondary antibodies at room temperature, (6) washed 3 times like in step 4, and (7) mounted in an anti-fade medium already containing Hoechst (Invitrogen #P36981). All of the following primary and secondary antibodies were diluted in 1XPBS containing 5% donkey serum. To establish our cerebral organoid cultures were differentiated into early neural tissue, we co-stained all cells with a mouse anti-NESTIN antibody (Santa Cruz #SC23927) and rabbit anti-PAX6 antibody (Biolegend #901301) using a 1/50 dilution for each. To visualize antigens of interest, cells were co-stained with a donkey anti-rabbit-AlexaFluor488 (Invitrogen #A32790) and donkey anti-mouse-AlexaFluor555 (Invitrogen #A32773) antibody using each at a dilution of 1/500. Lastly, cells were imaged at 63X on a Zeiss LSM 880 confocal laser scanning microscope.

### Protein extraction and quantification

To individually isolate individual nuclear and cytoplasmic protein fractions from iPSCs, subcellular fractionation was performed on live iPSCs using a commercially available kit (Thermo Scientific #78833) containing a 1X Protease and Phosphatase Inhibitor Cocktail (Thermo Scientific #78442) per manufacturer instructions. All protein isolates were subsequently handled on ice and stored at −80C. Prior to western blotting, the concentration of each protein fraction was quantified in duplicate using a Pierce BCA protein assay kit (Thermo Scientific #23225) and results were read on a Synergy H1 Microplate Reader using BioTek Gen5-3.04 software per manufacturer instructions.

### Fluorescent Western Blotting and ImageJ quantification

Protein extractions were freshly prepared under denaturing conditions by first diluting samples in 4x Laemmli Sample Buffer (Biorad #1610747) containing 50mM β-mercaptoethanol (BME; Sigma #M-7522) and then boiling diluted samples for 10 minutes at 100C in a T100 Thermal Cycler (Biorad #1861096). After denaturing, 8.5 ug of each protein sample was loaded onto a 4–20% Stain-free Protein Gel (Biorad #5678093) with a dual-fluorescent ladder (Biorad #1610374). Samples were then run in gel electrophoresis system (Biorad #1656019) using 1X Tris-Glycine-SDS running buffer (Fisher Scientific #BP13414) for 70 to 80 minutes at 130 V. Proteins were then transferred onto nitrocellulose membranes (Biorad #1704271) using a semi-dry blotting transfer system (Biorad #1704150). Membranes were then blocked at room temperature for 1 hour with gentle agitation in a blocking buffer made of: 5% non-fat milk (Nestlé #12428935), 0.1% Tween20 (Fisher Scientific #BP337-100), and 1X Tris-Buffered Saline (TBS; Boston Bioproducts #BM3014L). After blocking, all primary antibodies were diluted in this blocking buffer and allowed to incubate with membranes at 4C overnight with gentle agitation. Specifically, membranes were incubated with the following primary antibodies at the indicated dilution: mouse anti-KAT6A (Santa Cruz #sc293283 at 1/500), rabbit anti-KAT6B (Abcam #ab246879 at 1/1000), rabbit anti-NANOG (Cell Signaling #4903S at 1/1000), and mouse anti-OCT3/4 (Cell Signaling #75463 at 1/1000). The following day, membranes were washed 3 times at room temperature in 1X TBS supplemented with 0.1% Tween20 (TBST) with gentle agitation for 10 minutes each wash to remove excess antibodies. Membranes were then incubated for 1 hour at room temperature with secondary antibody solutions using gentle agitation. Similar to primary antibody solutions, all secondary antibodies were diluted in blocking buffer: goat anti-mouse-IR-680 (LI-COR #92668070 at 1/5000), goat anti-rabbit-IR-800 (LI-COR #92632211 at 1/5000), and anti-Tubulin-rhodamine (Biorad #12004165 at 1/1000). Next, membranes were washed 3 times at room temperature in TBST with gentle agitation for 10 minutes each wash to remove excess antibodies. Lastly, the fluorescent signals from the resulting membranes was visualized on a ChemiDoc Imaging System with a black tray (Biorad #12003028) using multiplexed-fluorescent imaging settings.

To quantify the changes in relative protein abundance of each antigen of interest from the fluorescent western blots, we imported the blot images from each channel (i.e. IRdye680, IRdye800, and Rhodamine) into a highly-cited open source imaging software analysis called ImageJ ^173^. Once each image was imported into imageJ, we selected each band of interest with the rectangular tool in ImageJ and ran a widely-used macros plugin to automatically quantify and normalize the regions of interest to the background. Specifically, we used the following setting with the Band/Peak Quantification macros: background width pixel set to 3, estimate background from top/bottom, and background estimation calculated from median. The macros’ algorithm then performs the same tasks as commercially available quantification software like Image Studio Lite from LI-COR Biosciences ^174^. Next, to determine the relative abundance of each antigen of interest, the resulting ‘signal’ for the antigen of interest was then normalized to the ‘signal’ for each respective loading control band. Lastly, to determine if the normalized antigen of interest signal was significantly different between the two conditions (ARTHS and control), we performed two-tailed student’s t-tests on the normalized antigen of interest signal in accordance with accepted statistical standards appropriate for each comparison^175,176^.

### RNA extraction, Quantification, and Quality Control

For whole transcriptome RNA-sequencing analysis, total RNA was extracted from all iPSC lines and cerebral organoids in triplicate. Total RNA was extracted from 1 million iPSCs that were snap-frozen on dry ice prior to extraction as previously described in the genotyping section. Specifically, total RNA was extracted from snap-frozen iPSC pellets using a PureLink RNA extraction kit (Invitrogen #12183018A) in conjunction with an on-column PureLink DNAse treatment (Invitrogen #12185-010). At day 15 and day 25 of cerebral organoid differentiation, total RNA was extracted from live cerebral organoids that were between 30uL to 50uL in volume upon pelleting in 1.5 mL polypropylene tubes (Thermo Scientific #3451) using a Qiagen RNA extraction kit (Qiagen #74004) with an on-column Qiagen DNAse treatment (Qiagen #79254). All DNA-free RNA was eluted in sterile nuclease-free water (Invitrogen #10977023) then stored at −80C.

Next, RNA extractions were quantified in duplicate using a Qubit fluorometer (Invitrogen #Q10211). Then to assess the RNA Integrity (RIN) prior to library preparation, 2uL of each RNA sample was analyzed on an Agilent 4150 Tapestation System using High Sensitivity RNA Screen Tapes (Agilent #5067-5579) and ladder (Agilent #5067-5581) per manufacturer instructions. RNA extractions with a RIN value less than 8.2 were not used to prepare subsequent RNA-sequencing libraries.

### RNA-sequencing: TruSeq Library Preparation, Quantification, Quality Control, and Sequencing

Total RNA was converted into DNA libraries using a TruSeq Stranded Total RNA Library Prep Gold kit (Illumina #20020599) and 96-well plates (Thermo Scientific #N8010560) which were sealed with microfilms (Thermo Scientific #4306311). Furthermore, all nucleic acids related to this protocol were stored at −80C and all the following bead-based clean up steps were performed using the following reagents: AMPure XP beads (Beckman Coulter #A63881), 200 proof ethanol (Fisher Scientific #BP2818500), sterile nuclease-free water (Invitrogen #10977023), and a 96-well magnetic stand (Invitrogen #AM10027).

First, to ensure all RNA samples were converted into RNA-sequencing libraries using identical inputs, all RNA samples were diluted to 93.3 ng/uL in ELB buffer (Illumina #20020599). Next, to simultaneously fragmented and depleted samples of ribosomal RNA (rRNA), 700 ng of diluted RNA was mixed with 8.5 uL of EPH buffer (Illumina #20020599) and 1 uL of FastSelect rRNA-depletion enzyme (Qiagen #335377). To complete the fragmentation and deletion process, this 17 uL reaction was then placed on a T100 Thermal Cycler (Biorad #1861096) using the following program recommended by the Qiagen FastSelect handbook: 94C for 8 minutes, 75C for 2 minutes, 70C for 2 minutes, 65C for 2 minutes, 60C for 2 minutes, 55C for 2 minutes, 37C for 2 minutes, 25C for 2 minutes, and then hold at 4C. The depleted and fragmented RNA was then converted into cDNA using random hexamers following the library preparation workflow described in Illumina’s TruSeq Stranded Total RNA Library Reference Guide (Document #1000000040499-v00). Specifically, the 17 uL reaction from the previous step was directly used to ‘synthesize first strand cDNA’ by combining this reaction with 8 uL of a master mix containing a ratio of 1 uL of Superscript Reverse Transcriptase II (Invitrogen #18064-014) to 9 uL of FSA (Illumina #20020599). After cDNA synthesis, the reactions were cleaned up using AMPure XP beads and then adenylated at the 3’ ends according to Illumina’s TruSeq Stranded Total RNA Library Reference Guide. Non-overlapping unique dual indexes (Illumina #20022371) were then added to each reaction and ligated according to manufacturer instructions. Following adapter ligation and AMPure XP bead clean up, the DNA libraries were PCR amplified using primers specific to fragments containing adapter sequences per manufacturer instructions. Enriched DNA fragments were then cleaned up using AMPure XP beads and final DNA libraries eluted in 30 uL of RSB (Illumina #20020599).

Before pooling the resulting double stranded libraries for next-generation sequencing, all libraries were quantified in duplicate using a Qubit fluorometer (Invitrogen #Q32854). Then the quality of each library was determined by analyzing 1uL of each sample on an Agilent 4150 Tapestation System using D1000 Screen Tapes (Agilent #5067-5582). To determine the average size of each library in base pairs (BP), this screen tape data was analyzed on TapeStation software to include all fragments ranging from 100 BP to 800 BP. Using the aforementioned library preparation protocol, each library ranged from 300 BP to 350 BP, with an average size of 320 BP. Next this data was used to calculate the molarity of each DNA library using the following formula:

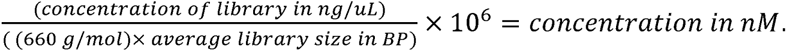

Lastly, all libraries were diluted to 10 nM in sterile nuclease-free water and then equivalent volumes of each 10nM library were combined to create a single equimolar pool that was submitted to UCLA’s Technology Center for Genomics and Bioinformatics (TCGB) core for next-generation sequencing; the resulting 10 nM pool was also analyzed using a Qubit fluorometer (Invitrogen #Q32854) and an Agilent 4150 Tapestation System (Agilent #5067-5582) as previously described to ensure individual libraries were correctly diluted and pooled. This resulting pool was sequenced on a 300-cycle S4 flow cell using an Illumina Novaseq6000 instrument for an average coverage of 77.1 million paired-end reads per sample across all 36 RNA-seq libraries and we did not note a substantial difference in coverage across conditions.

### Long Read RNA-sequencing: Iso-seq Library Preparation, Quantification, Quality Control, and Sequencing

Total RNA was shipped on dry ice to the University of Washington (UW) PacBio Sequencing core facility for PacBio’s full-length isoform sequencing (Iso-Seq). At UW, total RNA was converted into cDNA using a Iso-Seq Express cDNA generation kit (PacBio #101-737-500). UW then prepared Iso-seq libraries from this cDNA using a Iso-Seq Express SMRTbell library prep kit (PacBio #100-938-900). The resulting iso-seq libraries were then sequenced by UW on a PacBio Sequel II system using an 8M SMRT cell (v2.1 chemistry). The Sequel II system generated a total of ∼4.7 million HiFi reads which translated to a total of ∼3.6 million high quality full length non-chimeric (FLNC) circular consensus sequencing (CCS) reads, with an average of ∼453,000 FLNC reads per sample. The UW processed and analyzed the raw data using SMRT link software (version 10.1) to identify FLNC reads.

### RNA-sequencing: Differential Gene Expression and Principal Component analyses using DESeq2

After NGS by UCLA’s TCGB core, RNA-seq data was demultiplexed and fastq sequencing files were generated for all samples. Briefly, RNA-seq data was processed using the Arboleda lab’s RNA-seq pipeline as follows. First, the quality of fastq files was assessed using FastQC (v0.11.8)^177^. Next, fastq files were mapped to the human genome by aligning raw reads against the Gencode human genome (v31, GRCh38)^178^ using STAR (v2.7.0e) to generate bam files ^179^. Gene expression was then estimated by generating gene count matrices from the STAR-mapped raw reads and a GTF file of the human gene annotations from Gencode’s primary assembly (v31, GRCh38) using FeatureCounts (v1.6.5)^180^ from the Subread package ^181^. In the gene count matrices, only reads that uniquely mapped to exons were counted for each gene.

We performed principal component (PC) analysis as follows. A blinded variance stabilizing transformation (VST) was applied to raw read counts from gene count matrices using DESeq2 (v1.32.0) ^92,93^— followed by PC analysis and visualization with PCAtools (v2.4.0) ^182^. Then to quantify changes in gene expression observed in ARTHS samples relative to control samples in terms of log_2_FC(ARTHS/control), differential gene expression (DGE) analysis was performed on gene count matrices using DESeq2 which internally normalize for library size and uses a negative binomial distribution to model the data. Hypothesis testing in the DGE analysis was performed with the Wald’s test of statistical significance. Genes with a Benjamini and Hochberg adjusted p-value (P_adj_) less than 0.05 were classified significantly differentially expressed (sigDE) in ARTHS relative to controls, and log_2_FCs were shrunk using the DESeq2 ‘lfcShrink()’ function set to “apeglm” for approximate posterior estimation of GLM coefficients ^183^. These lists of sigDE genes were then used in downstream analyses for biological interpretation through various bioinformatics analyses described in other methods sections.

### RNA-sequencing: Enrichment analyses using ClusterProfiler and HOMER

Enrichment analyses were performed on DESeq2-identified lists of genes which were sigDE in ARTHS RNA-seq data, relative to controls, with ClusterProfiler (v4.4.4) ^94,184^ and <u>H</u>ypergeometric <u>O</u>ptimization of <u>M</u>otif <u>E</u>n<u>R</u>ichment (HOMER, v4.9) ^94,184^. Gene ontology (GO) enrichment tests were performed with the ClusterProfiler “enrichGO()” function across 3 categories (BPs, CCs, MFs) by submitting each list of sigDE genes against a background list comprised of all the human genes in Gencode (v31, GRCh38). GO terms were classified as significantly overrepresented (i.e. enriched) upon achieving both a Benjamini-Hochberg adjusted p-value less than 0.05 and a q-value less than 0.05 in each test. Hypergeometric motif enrichment tests were performed using HOMER’s “findMotifs.pl” command with default settings across 2 motif categories (de novo, known) and 3 motif lengths (8, 10, 12) to identify significantly overrepresented motifs (p-value<0.05) within the promoter regions of sigDE gene. Specifically, each list of sigDE genes was compared against HOMER’s default ‘human promoter set’ which is constructed from human Reference Sequence (RefSeq) database genes.

### RNA-sequencing: Partitioned Cell type, RNA-Binding Protein Target, and Disease Enrichment Analyses. Cell type enrichment

We used cell type marker genes of mid-gestation human neocortex as previously defined ^42^. Briefly, marker genes were identified as those differentially expressed in one cluster as compared to all other cell clusters in the dataset, and which are expressed in at least 10% of the cells of the cluster. Enrichment of cell type markers with sigDE ARTHS CO.15 and CO.25 gene lists was performed using Fisher’s exact test followed by FDR-correction (adjusted P<0.05). The background gene set used was all genes detected in the human neocortex dataset from which the cell type markers were derived ^42^.

### RBP target enrichment

For the set of brain-enriched RBP targets, we downloaded supplemental files of genes identified as containing splicing (AS) or gene expression (GEX) changes after RBP knockdown or knockout and those found through direct binding (CLIP). Relevant brain-enriched RBP target datasets were chosen using the following criteria – a) the RBP is expressed predominantly in neurons, or is differentially expressed between neural progenitors and neurons, b) knockdown/knockout or binding of the RBP was assayed in the relevant cell type (e.g. Ptbp1 knockout in neural stem cells), c) the RBP is expressed in mid-gestation human neocortex.

For the ENCODE eCLIP datasets (www.encodeproject.org), we chose all RBPs that a) are expressed in mid-gestation human neocortex, b) were assayed in the HepG2 liver cell line and c) have at least two replicate samples. See **Table S18** for citations associated with each public dataset and metadata.

When applicable, we filtered target lists using quality-control or p-value criteria provided in the publication. For datasets generated from mouse tissue, we retained genes with one-to-one human-to-mouse homology. For ENCODE datasets, we downloaded unfiltered eCLIP bed files for the RBPs described in [Van Nostrand et al.] and applied filtering criteria as described in the methods (peaks with P<0.001 and log2 fold-change ≥3 kept). Direct binding datasets (CLIP for brain-enriched or eCLIP for ENCODE) that were provided as genomic coordinates were overlapped with known gene coordinates using the GRanges R package and relevant genomic annotation. For datasets generated in mice, we used mm9 or mm10 depending on the dataset, and for human datasets we used hg38. We required that the coordinates for each (e)CLIP peak have at least 50% overlap with gene coordinates to assign it to that gene.

After QC and filtering steps, we performed enrichment analyses of the overlap between each RBP target set and sigDE ARTHS IPSC, CO15 and CO25 gene lists. Enrichment was performed using a one-sided Fisher’s exact test followed by FDR-correction for multiple testing (adjusted P<0.05). We used the union of genes detected in short-read IPSC, CO15 and CO25 sequencing as the background gene set, except for overlaps using datasets generated in mice, where we required genes in the above background to have one-to-one homology.

### Rare variant enrichment

We conducted enrichment analysis to localize rare-variant association signals from large-scale whole exome and genome sequencing studies of neurodevelopmental and psychiatric disorders within sigDE ARTHS genes merged across all timepoints. Logistic regression controlling for gene length was used to assess enrichment. We used all protein-coding genes as the background and set the significance threshold to an FDR-corrected pvalue<0.05 across all tested features. The gene list for ASD.SFARI.1or2 was compiled according to the indicated SFARI gene scoring (https://gene.sfari.org/database/gene-scoring/). The manually curated ‘Epilepsy.helbig’ list was downloaded from: http://epilepsygenetics.net/wp-content/uploads/2023/01/Channelopathist_genes_internal_2023_v2.xlsx

### RNA-sequencing: Autism Spectrum Disorder Risk Gene Analysis on unpartitioned sigDEGs

We performed an overlap analysis between the SFARI ASDR gene list and the DESeq2-identified lists of genes which were sigDE in ARTHS RNA-seq data relative to controls. In this ASDR gene overlap analysis, we first filtered all the genes that were tested in the DESeq2 analysis based on sufficient read coverage for statistical testing, then these expressed genes were compared to SFARI’s ASDR gene list to generate a final list of 1,066 ASDR genes that was then used in Fisher’s exact testing. Using SFARI’s scoring, we identified a subset of 366 ASDR genes which we determined ‘high confidence’ that had a gene scoring of 1 or S. We assessed the significance of the overlap between genes sigDE in ARTHS and ASDR genes from the full SFARI list (n= 1,066) and the high confidence ASDR gene list (n= 366) with Fisher’s exact tests.

We then constructed ASDR gene protein-protein interaction analysis with STRING. The 98 high-confidence ASD risk genes which were sigDE in ARTHS were also analyzed using the STRING database (version 11.5), to generate connections between proteins of interest based on physical and functional annotations from other databases. The resulting protein-protein interaction network was generated with a minimum required interaction score of 0.4, with the thickness of the lines reflecting the strength of the data support for a given interaction. The output from STRING was then visualized using Cytoscape (version 3.9.1). We then determined which Monarch Human Phenotype Ontology (HPO) terms were significantly overrepresented amongst these 98 genes relative to the number expected from a random network of the same size. In addition to the metrics produced by STRING’s enrichment algorithm, denoted by a ‘STRING’ prefix, odds ratios were calculated using Fisher’s Exact Tests based on the total number of human proteins available in STRING as of January 2023 (19,566) and using a Benjamini-Hochberg procedure to correct *p*-values for multiple testing of the current total number of Monarch HPO terms (10,634). Lastly, the nodes were colored based on selected HPO terms and a subset of the full network was generated to showcase connections between ARTHS phenotypes and ASD genes.

### Long Read RNA-sequencing: Autism Spectrum Disorder Risk Gene Analyses

Starting from FLNC, clustered Iso-Seq reads provided by UW, we performed alignment to GRCh38 with minimap2 (v2.24) ^185^. We then collapsed redundant isoforms using the ‘collapse_isoforms_by_sam.py’ script from the Cupcake collection of Iso-Seq post-processing scripts (v28.0.0, https://github.com/Magdoll/cDNA_Cupcake). We used additional Cupcake scripts to gather full length counts across samples. Taking the collapsed isoforms GTF file and full length counts table from Cupcake, we ran SQANTI3 (v4.2) to classify our isoforms according to novelty against the GENCODE 31 reference annotation and assess their quality ^186^.

Next, we prepared a transcriptome index using Salmon (v1.8.0) ^187^. To account for the presence of both previously annotated and novel transcripts in the short-read RNA-Seq data, we first used gffcompare (v0.12.6) to merge the GENCODE 31 reference annotation with our own annotation containing novel transcripts ^188^. We then created a FASTA file containing the nucleotide sequences of our merged annotation and the genome sequences of GRCh38 as a decoy to improve mapping performance. We ran the ‘salmon index’ command on our FASTA file, specifying the decoys and ‘-k 31’. We then quantified the short-read RNA-Seq FASTQ files for the IPSC and CO25 samples against our transcriptome index, using the ‘salmon quant’ command to run Salmon in mapping-based mode (with flags ‘-l A --validateMappings --seqBias --gcBias’).

Finally, we used the IsoformSwitchAnalyzeR package (v1.16.0) to detect differential transcript usage (DTU) between ARTHS and control samples ^189–192^. We imported our quantifications from Salmon using the ‘importIsoformExpression()’ function and created a design matrix that specified each sample as either ARTHS or control and also included stage of differentiation (iPSC or organoid) as a covariate. We then used the ‘importRdata()’ function to import our GTF and FASTA files mentioned above, and finally the ‘isoformSwitchTestDEXSeq()’ function to perform DTU analysis using DEXSeq. In these isoform-specific analyses, we controlled for the stage of CO differentiation in the DIU and DIE analysis to define how pathogenic KAT6A mutations affect transcription of ASDR genes irrespective of the differentiation. To determine the effect size of DIU between ARTHS and controls, the software first quantifies the fraction of gene expression originating from each observed transcript (i.e. isoform) assembly, which is called the isoform fraction (IF), and then it calculates the difference in IF (dIF) between the two conditions.

### Long RNA-sequencing: RNA-Binding Protein Target Enrichment Analyses

We performed enrichment analyses of the overlap between RBP target sets and genes with significant ARTHS DIU at the CO.25 time point as described above. Genes with significant DIU were determined as those having a gene_switch_q_value <0.05.

## Author Contributions

A.A.N. and V.A.A. conceptualized the project and designed all experimental approaches. A.A.N. and V.A.A. wrote and edited the manuscript with input from all authors. A.A.N. performed all experiments, curated all data—in addition to supervising and managing all components of this study. A.A.N. designed and executed all computational analyses related to the RNA-seq data. C.T.J. and M.J.G. analyzed the IsoSeq data and performed the isoform-centric DIU/DIE analyses. C.K.V. and L.T.U performed the cell type, RBP-related, ASD/Epilepsy-related gene enrichment analysis on partitioned RNA-seq and long read RNA-seq data. S.L.N.J. performed the ASD/HPO-related gene enrichment analysis on unpartitioned RNA-seq data. L.B. performed the QuPath analysis. C.J.O. assisted in the immunofluorescence experiments under the guidance of A.A.N.

## Acknowledgements

We would like to acknowledge the following core facilities for providing their technical expertise in helping generate some of the data from this manuscript: the Technology Center for Genomics and Bioinformatics (TCGB) at UCLA sequenced the short-read RNA-seq libraries generated by A.A.N., the PacBio Sequencing Services at University of Washington created and sequenced the ISO-seq libraries, and the Translational Pathology Core Laboratory (TPCL) at UCLA performed the histological processing of the cerebral organoids. We would also like to thank Angela Wei for establishing computational resources in the Arboleda lab that aided in some of the RNA-seq analyses. We would like to thank the families and the KAT6A foundation for their encouragement and support of this research.

## Abbreviations

NPC: neuroepithelial stem/progenitor cell
ND: neurodevelopment
ASD: Autism Spectrum Disorder
ARTHS: Arboleda-Tham syndrome
KAT: lysine acetyltransferase
IPSC: induced pluripotent stem cell
CO: cerebral organoid
CO15: cerebral organoids at day 15 of differentiation
CO25: cerebral organoids at day 25 of differentiation
RNA-seq: RNA sequencing
srRNA-seq: short-read RNA sequencing
Iso-seq: PacBio’s long-read RNA isoform-sequencing (Sso-seq) technology
sigDE: significantly differentially expressed in ARTHS relative to unaffected controls
DIU: Differential Isoform Usage
DIE: Differential Isoform Expression
dIF = dIF: difference in Isoform Fraction
L2FC: Log2 of the gene/isoform expression change between ARTHS/control
OR: Odds Ratio
GO: gene ontology
GEO: gene enrichment ontology
BPs: Biological Processes
CCs: Cellular Components
MFs: Molecular Functions
ASDR: Autism Spectrum Disorder Risk
RBP: RNA Binding Protein
Mic: Microglia
Per: Pericyte
End: Endothelial
OPC: Oligodendrocyte Precursors
InCGE: Interneuron Caudal Ganglionic Eminence
InMGE: Interneuron Medial Ganglionic Eminence
ExDp2: Excitatory Deep layer 2 neurons
ExDp1: Excitatory Deep layer 1 neurons
ExM-U: Maturing Excitatory-Upper Layer-enriched neurons
ExM: Maturing Excitatory neurons
ExN: Newborn Migrating Excitatory neurons
IP: Intermediate Progenitor
PgG2M: Cycling NPCs - G2/M phase
PgS: Cycling NPCs - S phase
oRG: Outer Radial Glia
vRG: Ventricular Radial Glia

